# Functional effects of chimeric antigen receptor co-receptor signaling domains in human Tregs

**DOI:** 10.1101/749721

**Authors:** Nicholas A.J. Dawson, Isaac Rosado-Sánchez, German E. Novakovsky, Vivian C.W. Fung, Qing Huang, Emma McIver, Grace Sun, Jana Gillies, Madeleine Speck, Paul C. Orban, Majid Mojibian, Megan K Levings

## Abstract

Antigen-specific regulatory T cells (Tregs) engineered with chimeric antigen receptor (CARs) are a potent immunosuppressive cellular therapy in multiple disease models. To date the majority of CAR Treg studies employed second generation CARs, encoding a CD28 or 4-1BB co-receptor signaling domain and CD3ζ, but it was not known if this CAR design was optimal for Tregs. Using an HLA-A2-specific CAR platform and human Tregs, we compared ten CARs with different co-receptor signaling domains and systematically tested their function. Tregs expressing a CAR encoding wild-type CD28 were markedly superior to all other CARs tested in an *in vivo* model of graft-versus-host disease. In vitro assays revealed stable expression of Helios and ability to suppress CD80 expression on DCs as key *in vitro* predictors of *in vivo* function. This comprehensive study of CAR signaling-domain variants in Tregs can be leveraged to optimize CAR design for use in antigen-specific Treg therapy.

## Introduction

Adoptive cell therapy with regulatory T cells (Tregs) shows promise in prevention or treatment of undesired immune responses, with several clinical trials completed or ongoing (reviewed in (Gliwinski et al., 2017)). To date, most of these trials have used polyclonal Tregs with unknown antigen (Ag) specificity, but work in mouse models has shown that Ag-specific Tregs are significantly more potent in a variety of disease contexts (Hoeppli et al., 2016). We and others developed an approach to generate Ag-specific Tregs with known specificity using chimeric antigen receptors (CARs) which combine extracellular antigen binding domains and intracellular signaling domains (Boardman and Levings, 2019). In comparison to polyclonal cells, the resulting CAR Tregs have enhanced potency, as demonstrated in a variety of models, including colitis (Blat et al., 2014; Elinav et al., 2009; Elinav et al., 2008), experimental autoimmune encephalomyelitis (Fransson et al., 2012), transplantation (Boardman et al., 2017; Dawson et al., 2019; MacDonald et al., 2016; Noyan et al., 2017; Pierini et al., 2017), and immunity to therapeutic proteins (Kim et al., 2015b; Yoon et al., 2017).

The majority of CAR Treg studies to date have used the so-called “second generation” CAR design, which includes a single membrane-proximal co-stimulatory domain, followed by CD3ζ. Due to the extensive evidence for the important role of CD28 co-stimulation in Tregs (Esensten et al., 2016), the majority of CAR-Treg studies selected this protein as the source of co-stimulation. However, as with conventional T cells (Tconvs), Tregs express a number of co-receptors that provide co-stimulatory or co-inhibitory signals (reviewed in (Kumar et al., 2018)), the functional relevance of which may differ from that in Tconvs, and vary depending on the tissue/disease context. An additional consideration is that the vast majority of studies which have defined co-receptor function in Tregs used genetic-deletion models in which co-receptors of interest were absent throughout Treg development, thus making it difficult to infer function in the fully differentiated cells which would be used for CAR engineering.

Extensive research in the context of oncology has sought to optimize CAR design to deliver potent and persistent Tconvs (Sadelain et al., 2013). Much of this work focused on comparing co-receptor signaling domains, revealing a key role for co-stimulation in CAR function. For example, comparisons of CD28- to 4-1BB-based second generation CARs in CD8^+^ T cells, revealed that 4-1BB-based CAR T cells are more persistent and resistant to exhaustion, and express a memory-T-cell like pattern of gene expression (Long et al., 2015; Song et al., 2011) whereas CD28-based CAR T cells have a more acute anti-tumor effect (reviewed in (van der Stegen et al., 2015)). These findings led to intense research to define the optimal contexts in which to use 4-1BB-versus CD28-based CAR T cells, with both versions now in clinical use with Kymriah^®^ encoding CD28 and Yescarta^®^ 4-1BB (Salmikangas et al., 2018).

The functional effects of CARs encoding other co-receptor domains have also been examined in Tconvs. For example, in comparison to CD28 and 4-1BB, expression of an ICOS-based CAR results in higher secretion of Th17-associated cytokines and longer in vivo persistence in a xenograft tumor model (Guedan et al., 2014; Guedan et al., 2018). The effects of CARs encoding co-inhibitory receptors have also been explored. Expression of second-generation CARs encoding PD-1 or CTLA-4 together with CD28- or 4-1BB-encoding CARs is an effective approach to limiting toxicity by restricting off-target T cell stimulation (Fedorov et al., 2013). Thus, the function, potency and persistence of CAR-engineered Tconvs can be tailored by choice of co-stimulatory domain.

How Tregs would be affected by CARs encoding co-stimulatory domains other than CD28 or 4-1BB was unknown. Seeking to identify CARs that could modulate Tregs by altering their stability, cytokine production, survival and/or other properties of therapeutic benefit, we sought to comprehensively compare the *in vitro* and *in vivo* function of CAR-Tregs expressing second-generation CARs encoding one of 10 different co-stimulatory domains.

## Results

### Generation, cell surface expression and selection of signaling-domain CAR variants

We selected 9 co-receptor proteins to test in a CAR format in comparison to an existing CAR which encodes an intracellular CD28 domain and an extracellular single chain antibody (scFv) specific for HLA-A2, and stimulates potent Treg suppression in vitro and in vivo (Dawson et al., 2019; MacDonald et al., 2016). Co-receptors were selected from CD28 or TNFR family proteins (**Figure 1A**) since these are functionally relevant in T cells and, in some cases, had already been successfully used as CARs in Tconvs. Within the CD28 family, in addition to the wild type (wt) CD28 protein, which is known to be important for Treg activation and proliferation (Golovina et al., 2008; Tai et al., 2005; Zhang et al., 2013) we selected: CD28(Y173F), a point mutant with diminished PI3K pathway activity which may be beneficial for Treg function (Okkenhaug et al., 2001); ICOS, which is important in Treg survival and may be involved in IL-10 production (Kornete et al., 2012; Landuyt et al., 2019; Redpath et al., 2013); CTLA-4, which is essential for Treg function (Esensten et al., 2016; Walker and Sansom, 2011); CTLA-4 (Y165G), a point mutant with increases cell surface expression (Nakaseko et al., 1999); and PD-1, which is essential for generation and maintenance of peripherally-induced Tregs (Chen et al., 2014). Within the TNFR family, we selected: 4-1BB as this co-receptor is beneficial for the longevity of CAR-T cells (Long et al., 2015; Song et al., 2011) and in models of autoimmunity is beneficial for Tregs (Kumar et al., 2018); OX40 and GITR, which promote Tregs in certain contexts (Kumar et al., 2018); and TNFR2 which stimulates Treg proliferation and suppression (Chopra et al., 2016; Pierini et al., 2016).

**Figure 1.**
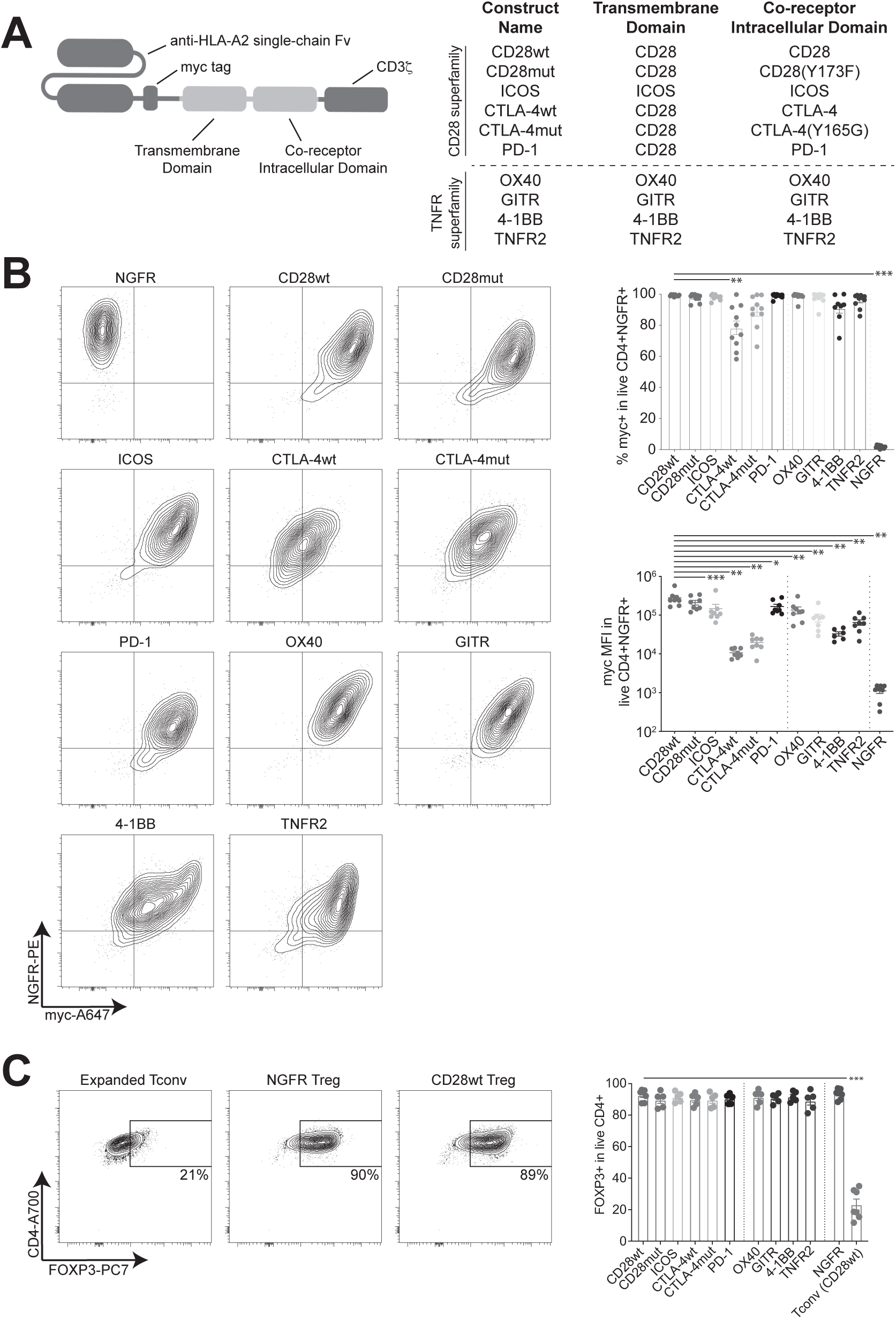
Design and surface expression of signaling domain CAR variants on human Tregs. **(A)** CARs consisting of an extracellular anti-HLA-A2 scFv, myc tag and CD8α stalk, followed by various transmembrane and co-receptor domains, and CD3ζ were cloned. The indicated constructs were selected on the basis of cell surface expression in transiently transfected 293T cells (Supplemental Figure 1). **(B)** CD4^+^CD25^hi^CD127^−^ human Tregs were sorted and transduced using lentivirus generated from the constructs in (A). After 7 days, transduced cells were purified on the basis of NGFR expression, and analyzed by flow cytometry. Shown are representative and averaged data depicted as the proportion or mean fluorescence intensity (MFI) of myc^+^ cells within live, CD4^+^ΔNGFR^+^ cells. n=6-8 from at least four independent experiments. **(C)** FOXP3 expression in CAR Tregs after 7-day expansion and purification. CAR Tregs were transduced, expanded, purified as in (B), then analyzed by flow cytometry for FOXP3 purity. Left: representative flow plots. Right: summarized data of percent FOXP3 expression within live CD4+ cells. n=5-6 donors, pooled from at least 3 independent experiments. Statistics in (B-C) show one-way ANOVA with a Holm-Sidak post-test to compare all constructs to CD28wt. Mean ± SEM, * p < 0.05, ** p < 0.01, *** p < 0.001.

Because the choice of transmembrane domain can affect CAR expression (Bridgeman et al., 2010; Dotti et al., 2014), two versions were created for several of constructs: one using the transmembrane domain from CD28 and the other using the “native” transmembrane domain from the co-receptor being tested. Sequences for the transmembrane and intracellular signaling portions of these proteins were taken from the UniProt database (Consortium, 2018), placed C-terminal to the anti-HLA-A2-specific scFv, a c-Myc epitope tag, and a CD8α stalk sequence. The full CAR construct was cloned into a bi-directional lentiviral vector encoding ΔNGFR as a transduction marker as described (MacDonald et al., 2016). Cell surface expression was first tested in 293T cells by quantifying the proportion and intensity of Myc-expressing cells within the ΔNGFR-positive population (**Supplemental Figure 1A**). Most CAR constructs had similar levels of surface expression with the CD28 or native transmembrane domains (PD-1, ICOS, OX40, 4-1BB, TNFR2), but the levels of surface expression of CTLA-4mut and GITR were significantly higher with the CD28 or native domain, respectively. To determine whether changing the scFv affected expression patterns, some of the co-stimulatory domains were also tested in the context of a CAR encoding an anti-HER2 scFv, revealing no significant differences in comparison to the HLA-A2-based CAR (**Supplemental Figure 1B**). On the basis of these data, with the exception of ICOS, the CD28 transmembrane domain was selected for all the CD28 family co-receptors, and the native transmembrane domain was selected for all the TNFR family co-receptors.

To test expression in human Tregs, CD25^hi^CD127^lo^CD4^+^ cells were sorted from peripheral blood, activated via their TCR and transduced with lentivirus encoding one of the 10 signaling domain variant A2-CARs, or an empty vector control. After 7 days, successfully transduced ΔNGFR^+^ cells were isolated and CAR expression was determined on the basis of the proportion and mean fluorescence intensity (MFI) of myc expression (**Figure 1B**). In terms of proportion, within the CD28 family, only the CTLA-4wt (but notably, not the CTLA-4mut) variant showed significantly lower A2-CAR expression than the CD28wt construct; within the TNFR2 series, there were no significant differences. In terms of MFI, differences in expression intensity were seen, with the CD28wt and CD28mut constructs consistently producing the highest levels of expression. No differences in FOXP3 expression were seen between the Treg groups post transduction and expansion (**Figure 1C**).

### Wild type CD28 signaling is required for optimal Treg suppression in vivo

To assess the function of the signaling domain A2-CAR variants with human Tregs, we used the xenogeneic graft-versus-host disease (GVHD) mouse model (Cooke et al., 1996; Hill et al., 1997) in which A2-CAR Tregs prevent GVHD via suppressed proliferation/engraftment of co-injected allogeneic HLA-A2^+^ PBMCs (MacDonald et al., 2016). CD25^hi^CD127^lo^CD4^+^ Tregs were sorted, stimulated, transduced and purified as described above, and expanded further by TCR-restimulation for 5 additional days. Immunodeficient NGS mice were irradiated, then injected with 10^6^ HLA-A2^+^ PBMCs in the absence or presence of 5×10^6^ (high ratio) or 2.5 x10^6^ (low ratio) of the indicated type of CAR Treg (**Figure 2A**). Mice receiving Tregs transduced with the CD28wt A2-CAR represented the positive control, and negative controls were mice injected with Tregs transduced with an antigen-irrelevant HER2-CAR, or a first-generation A2-specific CAR lacking a co-receptor signaling domain. Survival and GVHD score were monitored over time. As expected, mice which received CD28wt A2-CAR Tregs at either a high or low ratio were significantly more protected from GVHD than the negative control mice (**Figure 2B and Supplemental Figure 2A**).

**Figure 2.**
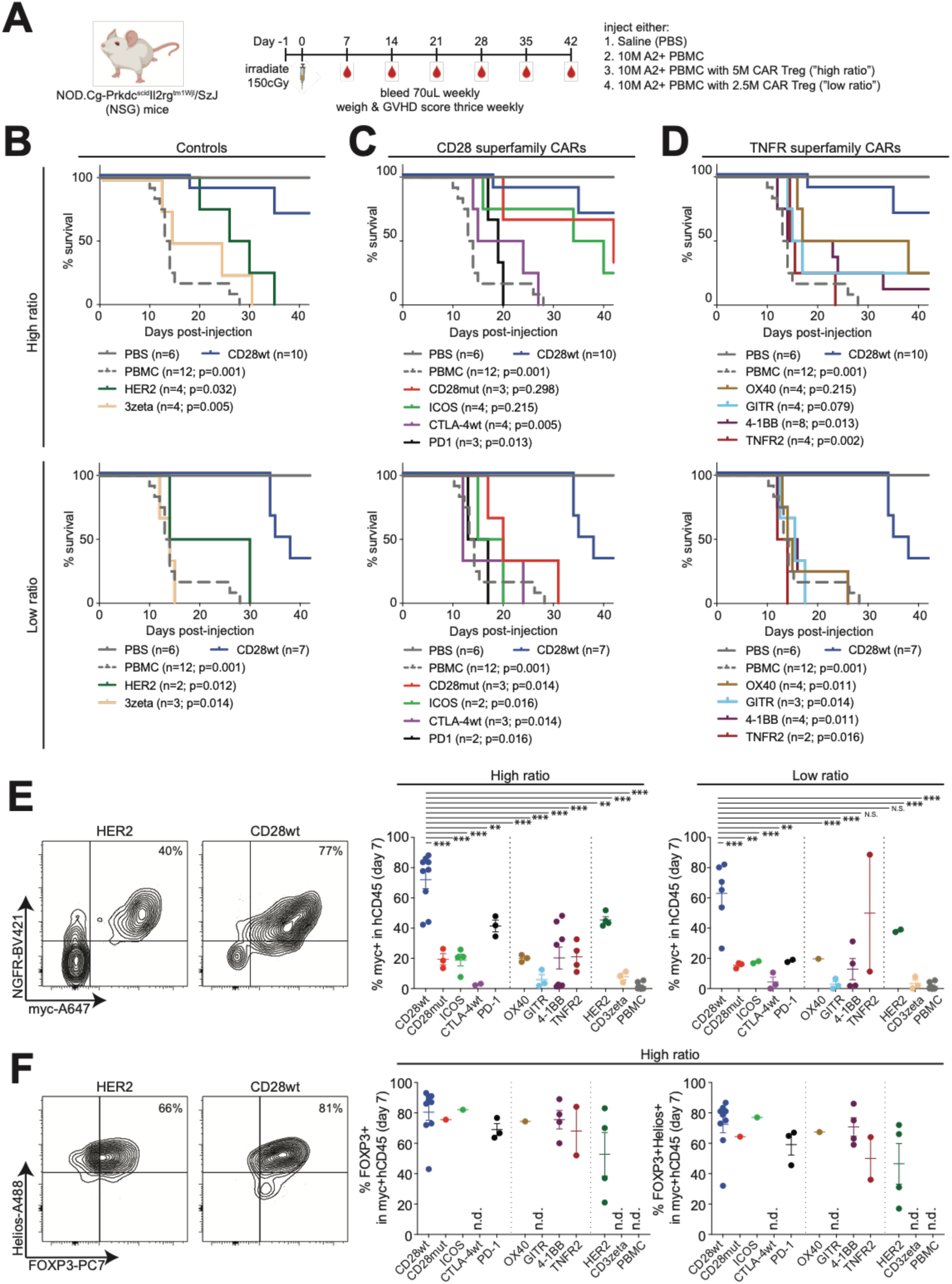
In vivo suppression of xenogeneic graft-versus-host disease by signaling domain CAR variant Tregs. 8 to 12-week old female NSG mice were irradiated one day before injection with PBS, or 10^7^ HLA-A2^+^ PBMC in the absence or presence of 2.5 or 5.0 x10^6^ of the indicated type of CAR-Treg. Mice were weighed and scored for GVHD thrice weekly and bled weekly for flow cytometry analysis. **(A)** Schematic design of the experiment. **(B-D)** Survival curves. **(B)** Positive (CD28wt) and negative (HER2, CD3zeta) controls. **(C)** CARs with signaling domains from the CD28 superfamily. **(D)** CARs with signaling domains from the TNFR superfamily. <u>Top:</u> Mice receiving a high ratio of Treg:PBMC (1:2). <u>Bottom:</u> Mice receiving a low ratio of Treg:PBMC (1:4). Data from PBS, PBMC and CD28wt Treg mice are repeated in each panel for comparison. Statistics show adjusted p values corrected for multiple comparisons for pair-wise log-rank Mantel-Cox tests, comparing the survival curve for all constructs to CD28wt. **(E-F)** Seven days post-cell injection the proportion of **(E)** CAR-expressing (myc^+^) cells within hCD45+ cells and, for mice with sufficient cells, **(F)** FOXP3^+^ (left) and FOXP3^+^Helios^+^ (right) cells within CAR-expressing (myc^+^) hCD45^+^ cells was determined. Representative and averaged data are shown. The number of individual mice in each group tested over three independent experiments are indicated in (B-D). Statistics show one-way ANOVA with Holm-Sidak post-test comparing all constructs with at least two data points to CD28wt. Mean ± SEM. * p < 0.05, ** p < 0.01, *** p < 0.001. “n.s.” denotes not significant. “n.d.” denotes no data where there were insufficient events to record a result.

Surprisingly, CD28wt A2-CAR Tregs provided better protection than all the other co-receptor CAR variants tested. At a high ratio, CARs encoding CD28mut or ICOS domains were not significantly different from the CD28wt construct but at low ratios, the CD28wt A2-CAR Tregs provided significantly better protection from GVHD (**Figure 2C**). At both high and low ratios, CARs encoding TNFR family co-receptors were ineffective in promoting CAR Treg function; at low ratios, survival and GVHD scores were even lower than in mice injected with irrelevant-antigen-specific HER2-CAR Tregs (**Figure 2D**). The effects on in vivo survival and GVHD scores were mirrored by engraftment and proliferation of human HLA-A2^+^ cells: mice which received CD28wt A2-CAR Tregs consistently had the slowest increase in human CD45^+^ cell engraftment (**Supplemental Figure 2B & 3**) and, in the high ratio mice, lower proportions of circulating HLA-A2^+^ cells at day 7 (data not shown).

We previously found that circulating CAR Tregs are difficult to detect >7 days after injection (Dawson et al., 2019). Since CARs encoding TNFR family domains can increase CAR T cell longevity (Long et al., 2015; Song et al., 2011), we sought to characterize CAR Treg engraftment and stability by analyzing the proportion of Myc^+^ cells within the hCD45^+^ gate over time. At day 7, there was large variation in the frequency (**Figure 2E**) and absolute number (**Supplemental Figure 4A**) of CAR Tregs between the constructs, but a clear and significant survival advantage for the CD28wt-CAR Tregs. Indeed, at day 14 only mice that received CD28wt-CAR Tregs (**Supplemental Figure 4B**) had detectable circulating cells.

Further characterization of CAR Tregs in mice in which they were detectable on day 7 after injection also revealed variation in the maintenance of the expected FOXP3^+^ phenotype, with the CD28wt-CAR Tregs having the most consistent and high proportion of FOXP3-expressing cells (**Figure 2F, Supplemental Figure 4C**). It has been reported that one property of CAR Tconv is their ability to acquire their target antigen through a process called trogocytosis (Hamieh et al., 2019). In all samples where Myc+ CAR T cells were detected, none were HLA-A2+, suggesting that at least in this model and at this time point, we were not able to detect trogocytosis (**Supplemental Figure 4D**).

### Signaling domain CAR variants differ in their ability to activate Tregs and stimulate cytokine production

The in vitro phenotypic and/or functional features of CAR Tregs that correlate/predict in vivo function are unknown. This series of signaling domain CAR variants provided an ideal opportunity to further explore this question as they had a range of effects in vivo, from highly effective (CD28wt), to moderately effective (e.g. CD28mut and ICOS) to ineffective (TNFR2). We first asked whether the CARs might differ in their ability to stimulate Treg activation. Signaling domain CAR-Treg variants were rested overnight, then stimulated with HLA-A2-expressing K562 cells, and after 24h, analyzed by flow cytometry for expression of the activation markers CD69 and CD71. In the absence of stimulation there were no significant differences in expression, demonstrating the absence of tonic signaling (**Supplemental Figure 5A).** There was a large variation in the capacity of the different CARs to activate Tregs, with CD28wt and CD28mut stimulating the highest, and PD-1 the lowest expression of both markers (**Figure 3A**). Importantly, the presence of each CAR variant did not affect the ability of CAR Tregs to be activated via the TCR (**Supplemental Figure 5A**).

**Figure 3.**
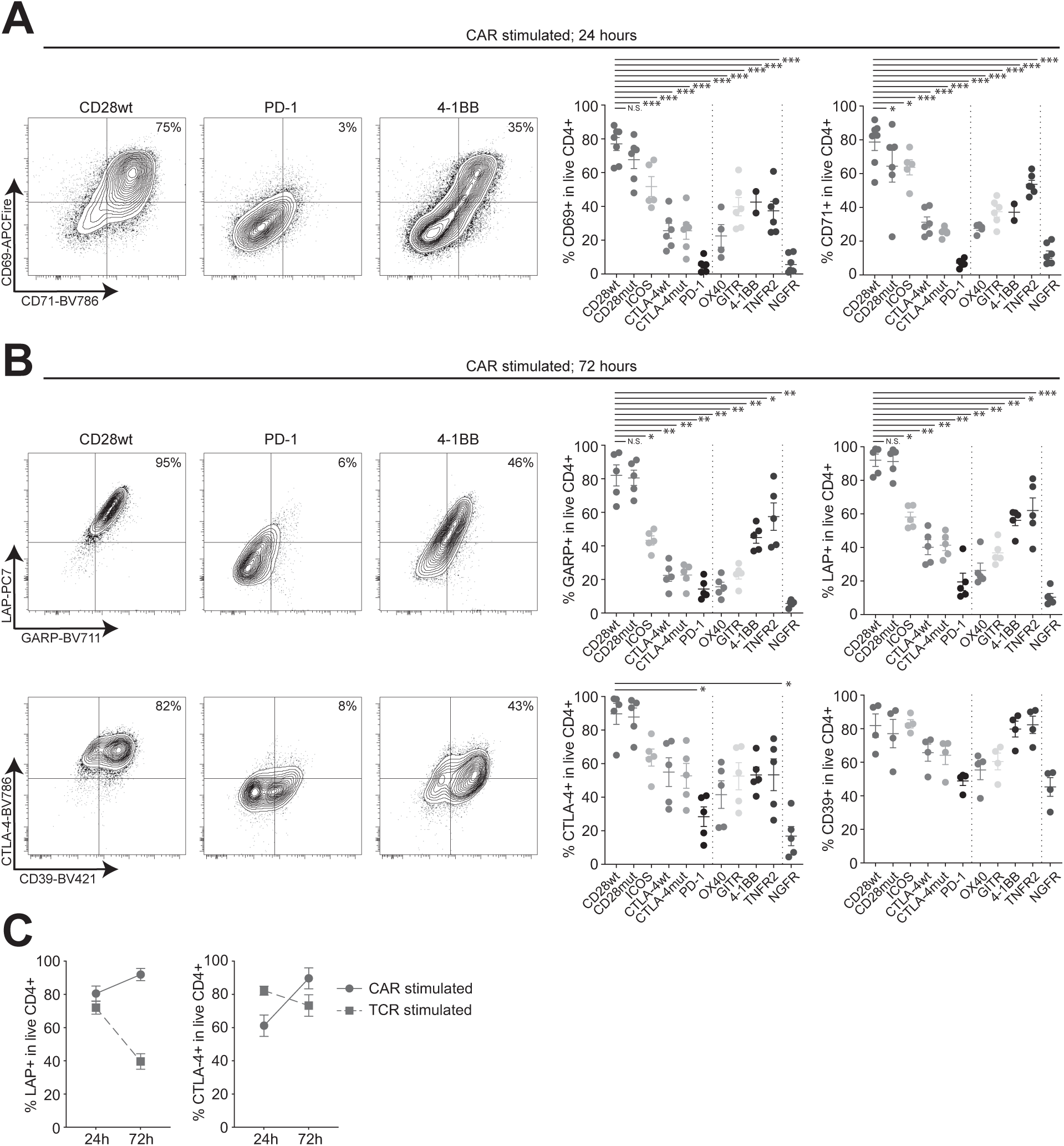
Signaling domain CAR variant-mediated expression of activation makers and suppressive proteins. Human Tregs were transduced and expanded, rested overnight in low IL-2 conditions, then co-cultured with irradiated K562 cells expressing HLA-A2 at a ratio of 1 K562 to 2 CAR Tregs. After 24 **(A)** or 72 **(B)** hours expression of the indicated protein was determined by flow cytometry. n=2-7 donors from a minimum of least two independent experiments. Statistics show one-way ANOVA with Holm-Sidak post-test comparing all constructs to CD28wt. **(C)** Kinetics of LAP^+^ (left) and CTLA-4^+^ (right) expression on CAR- or TCR-stimulated CD28wt-CAR Tregs from experiments in (A) and (B). Mean ± SEM. * p < 0.05, ** p < 0.01, *** p < 0.001. “n.s.” denotes not significant.

We next asked how the signaling domain CAR variants affected expression of key Treg-specific effector molecules, including LAP (inactive form of TGF-β), GARP (receptor for LAP), CD39 (ATP/ADP ectonucleoside) and CTLA-4. CAR Tregs were stimulated with HLA-A2-K562 cells for 24h (**Supplemental Figure 5B**) or 72h (**Figure 3B**) then analyzed by flow cytometry. Similar to results with CD69 and CD71, CD28wt- and CD28mut-CARs stimulated the highest expression of GARP, LAP and CTLA-4. Interestingly, several CARs that were non-functional in vivo (e.g. 4-1BB and TNFR2), exhibited high expression of CD39 upon CAR-stimulation, suggesting expression of this pathway does not correlate with in vivo function in the GVHD model. Comparisons between TCR- and CAR-stimulation revealed an interesting difference in kinetics: in response to CD28wt A2-CAR-, but not TCR-stimulation, expression of LAP and CTLA-4 was reinforced over time (**Figure 3C**).

To assess co-receptor domain-dependent variation in the cytokine profile of CAR-stimulated Tregs, supernatants were collected from cells stimulated via HLA-A2 for 72h and analyzed by cytometric bead assay. In comparison to Tconvs expressing a CD28wt CAR, all Tregs exhibited a low level of cytokine production, either when stimulated through the CAR (**Supplemental Figure 5C**) or TCR (data not shown). None of the CAR Tregs produced detectable amounts of IL-2, IL-4 or TNF-α when stimulated with A2 or the TCR. The CD28wt- and CD28mut-CAR Tregs did produce detectable levels of IFN-γ, IL-17A, IL-10 and IL-6 when stimulated via the CAR (but not the TCR), but the amounts were lower than those from the Tconv control. Notably, contrary to our prediction and a previous report in Tconvs (Guedan et al., 2014), the ICOS-encoding CAR did not stimulate production of IL-10 or IL-17A.

### 4-1BB- and TNFR2-encoding CARs destabilize Tregs

Since 4-1BB is an effective co-stimulatory domain in conventional CAR T cells (June and Sadelain, 2018; Lim and June, 2017), and TNFR2 signaling is well-known to positively affect Tregs (Chopra et al., 2016; Pierini et al., 2016), we were surprised to find that these two CARs not only did not perform as well as the CD28wt-CAR, but that they were also significantly worse than the irrelevant antigen-specific HER2-CAR control at protecting mice from GVHD (**Figure 2**). To further investigate the mechanisms underlying this finding, we carried out in vitro assays to more extensively interrogate the long-term effects of CAR-stimulated Treg activation. Tregs expressing one of the signaling domain variants, or control CARs, were stimulated with beads coated with HLA-A2, in the presence of 100U/mL IL-2. Cell number, expression of FOXP3 and Helios, and the amount of methylation in the Treg-specific demethylation region (TSDR) were monitored over 12 days (**Figure 4A**). Most CAR constructs did not stimulate Treg proliferation (**Figure 4B**): only CD28wt-, 4-1BB- and TNFR2-encoding CARs were able to stimulate a significant amount of proliferation in comparison to ΔNGFR control Tregs. Notably, the TNFR2-CAR stimulated significantly more proliferation than the 4-1BB- or CD28wt-CARs, with an average of >16-fold expansion over 12 days. Interestingly, this was the first in vitro assay in which we observed a clear difference between the CD28mut- and CD28wt-CARs, suggesting that this tyrosine residue (Y173) in CD28 is essential for stimulating Treg proliferation and that this may be an important property for in vivo function.

**Figure 4.**
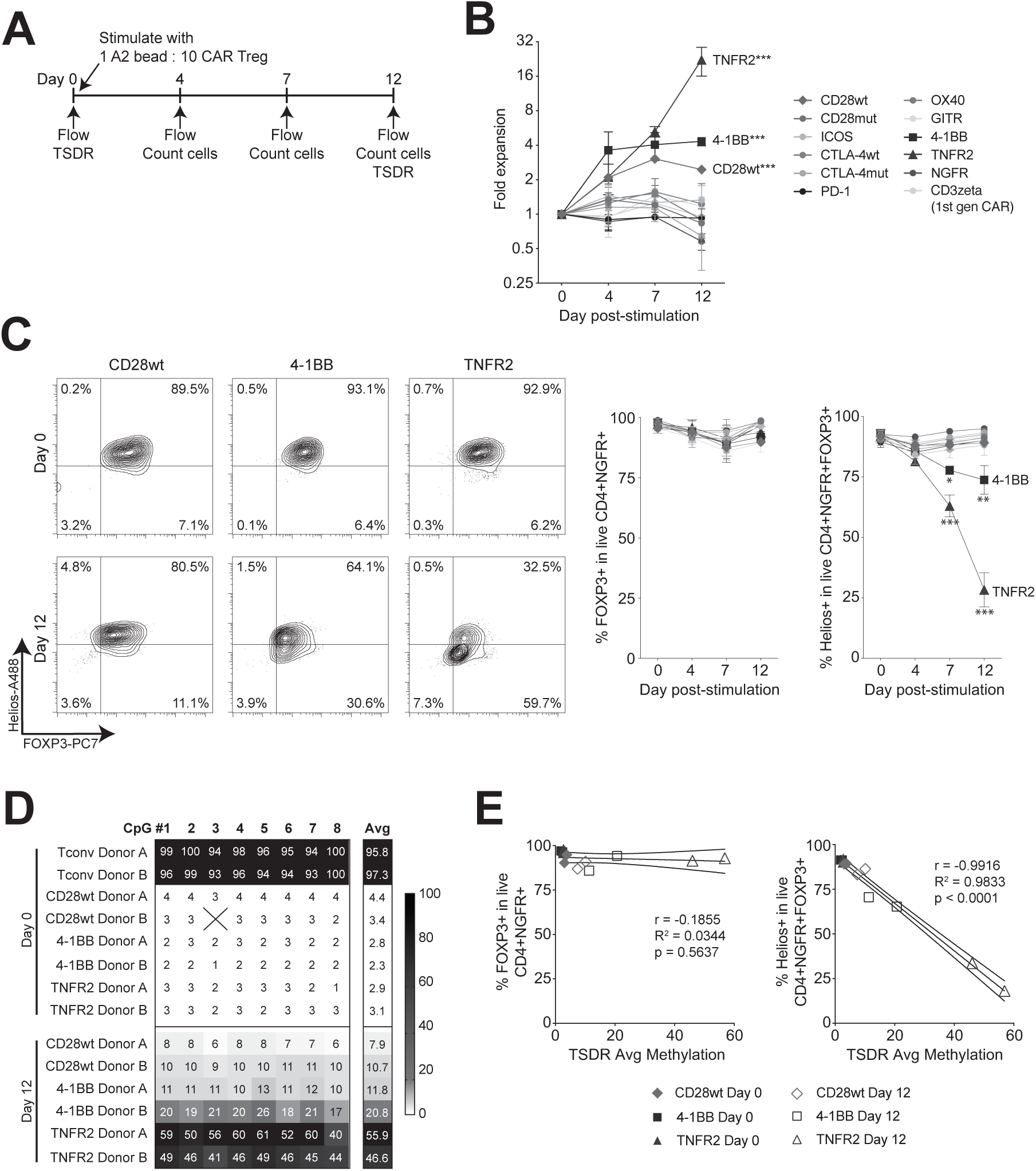
Effects of prolonged stimulation of signaling domain CAR variants on Treg stability. Naïve Tregs (CD4^+^CD25^+^CD127^−^CD45RA^+^) were sorted, transduced with the indicated signaling-domain CAR variants and expanded for 7 days. After resting overnight in low amounts of IL-2, CAR Tregs were stimulated with beads coated with HLA-A2 at a ratio of 1 bead to 10 CAR Tregs for 12 days. Cells were counted and analyzed by flow cytometry on day 4, 7 and 12. **(A)** Schematic diagram of experiment setup. **(B)** Cell expansion over time. Statistics show overall effects two-way ANOVA with Holm-Sidak post-test comparing all constructs to the NGFR group. **(C)** Flow cytometry analysis of FOXP3 and Helios within live CD4+NGFR+ cells before (day 0) or 12 days after bead stimulation. Representative and averaged data for proportion of FOXP3+ cells within live CD4+NGFR+ and Helios+ cells within live CD4+NGFR+FOXP3+ cells. Data for (A-C) are n=2-3 donors, pooled from at least two independent experiments. Statistics show two-way ANOVA with Holm-Sidak post-test comparing all constructs at each timepoint to CD28wt. **(D)** Pyrosequencing of cells lysed on day 0 and day 12 showing percent methylation of eight CpGs from CNS2 within the *FOXP3* locus, known as the Treg-specific demethylated region (TSDR). **(E)** Correlation of average TSDR methylation and percent FOXP3+ or percent Helios+ from C. For (D-E), n=2 male samples tested from one experiment. Pearson r statistic and two-tailed p value shown. Mean ± SEM. * p < 0.05, ** p < 0.01, *** p < 0.001.

Whereas analysis of FOXP3 expression showed no significant difference between the groups, Helios expression revealed significant differences which directly correlated with proliferation. Specifically, only 4-1BB- and TNFR2-encoding constructs lost significant amounts of Helios expression (**Figure 4C**). Since Helios expression is thought to be associated with Treg lineage stability (Kim et al., 2015a; Nakagawa et al., 2016; Skadow et al., 2019; Thornton et al., 2019) we next asked whether Tregs expressing 4-1BB- and TNFR2-encoding CAR constructs differed in their lineage stability. Accordingly, for these three constructs, we analyzed the methylation status of 8 CpG islands within the Treg-specific demethylated region (TSDR) (Baron et al., 2007; Polansky et al., 2008) of the *FOXP3* gene (**Figure 4D**). Before stimulation via the CAR (day 0), the 3 types of CAR Tregs did not differ in the amount of TSDR methylation, with an almost completely demethylated TSDR. However, after 12 days of stimulation with HLA-A2-coated beads, Tregs expressing the TNFR2-CAR had a marked, ~50% increase in methylation. Tregs expressing the 4-1BB-CAR exhibited an intermediate phenotype, trending toward a more highly methylated TSDR region than CD28wt-CAR expressing cells. Interestingly, we observed a strong negative correlation between TSDR methylation and expression of Helios, but not of FOXP3 (**Figure 4E**). These data suggest that loss of Helios expression is more highly correlated with Treg lineage instability than is loss of FOXP3.

We next asked whether differences in the proportions of naïve and memory cells might underlie the differential proliferative capacities of cells expressing the CD28wt, 4-1BB and TNFR2 signaling domain CARs, and those expressing the other CARs. The various CAR Tregs were stimulated via HLA-A2 for 7 days, then analyzed for expression of CCR7 and CD45RA in order to quantify naive, central memory, effector memory, and CD45RA^+^ effector memory cells. Most CARs, including the CD28wt, promoted expansion of Tregs with an effector memory phenotype, with the exception of the 4-1BB- and TNFR2-encoding CARs, which preserved high proportions of central memory cells, and the PD-1-encoding CAR, which enhanced the proportion of naive cells (**Supplemental Figure 6A**). By corollary, Tregs stimulated with 4-1BB and TNFR2 CARs also had a trend toward lower expression of PD-1, a cell surface molecule often inversely associated with cell expansion potential (**Supplemental Figure 6B**).

Overall, these data suggest that for function in vivo, CAR Tregs must be able to proliferate and acquire an effector memory phenotype, but that excessive and rapid proliferation is detrimental, leading to instability and an unfavorable central memory phenotype.

### Suppression of antigen-presenting cells is a better predictor of CAR Treg in vivo function than suppression of T cells

Tregs suppress many different types of immune cells via different mechanisms, with a growing appreciation for the important role of CTLA-4 mediated suppression of dendritic cells (Walker and Sansom, 2011) in addition to classical direct suppression of T cell proliferation. We next sought to measure the suppressive capacity of the signaling-domain CAR variants towards these two different cell types. To measure direct suppression of T cells, we set up an assay in which A2-negative T cells were pre-stimulated with anti-CD3/28 beads for 16h, the beads removed, and the cells were then cultured for 72h in the absence or presence of the various signaling-domain A2-CAR Tregs, which had been previously stimulated with A2-coated beads (**Figure 5A**). Consistent with the in vivo data (**Figure 2**), CD28wt-CAR Tregs suppressed both CD4^+^ and CD8^+^ T cells more potently than all other CAR Tregs (**Figure 5B-C**). However, divergently from the in vivo data, most CAR variants, including 4-1BB and TNFR2, were significantly better at suppressing T-cell proliferation than the control transduced ΔNGFR Tregs.

**Figure 5.**
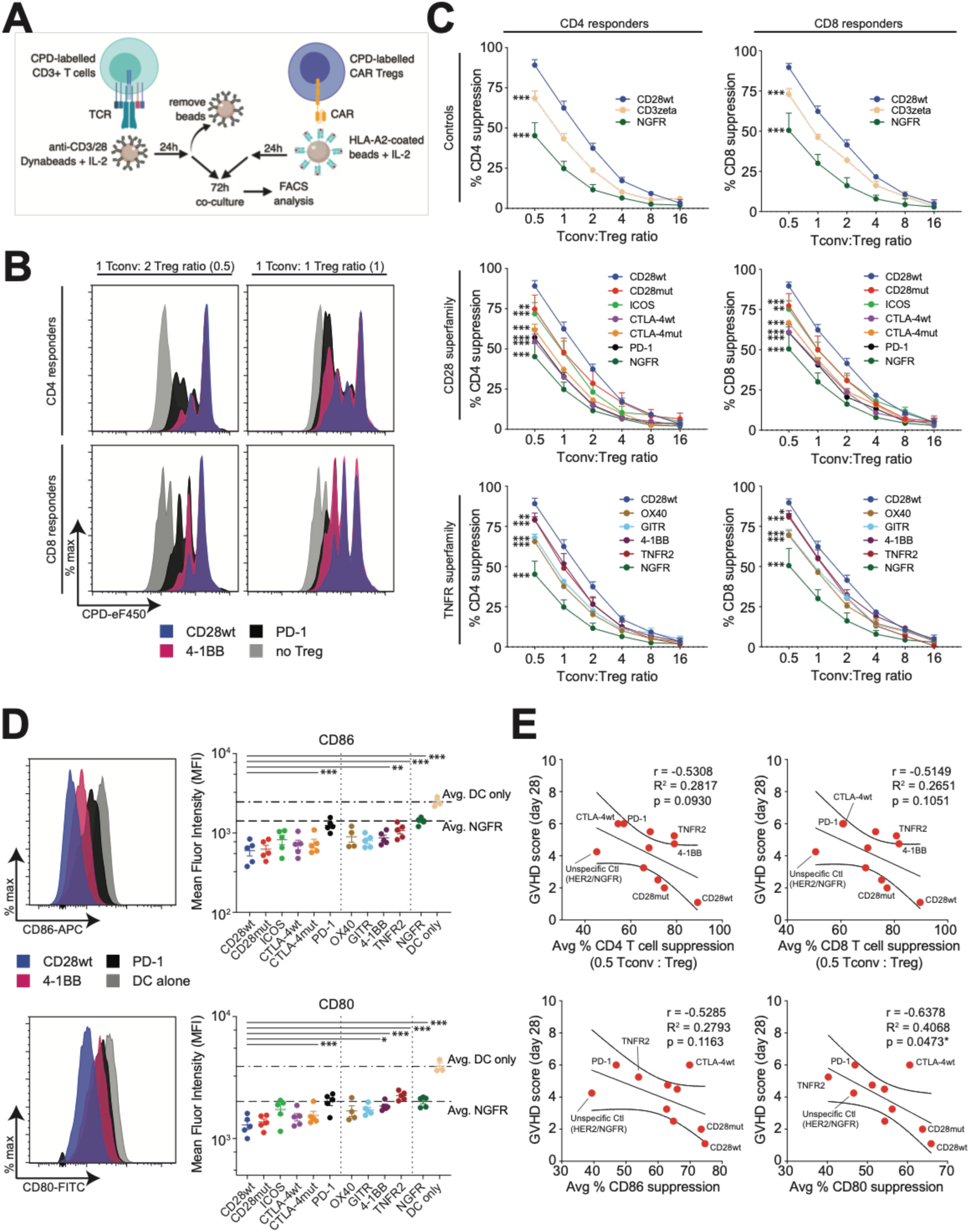
Suppressive properties of signaling domain CAR Treg variants. **(A-C)** In vitro indirect T cell suppression assay. Cell-proliferation dye (CPD)450-labelled responder HLA-A2^neg^CD3^+^ T cells were pre-stimulated for one day with anti-CD3/CD28 dynabeads and IL-2 before being co-cultured without or with different ratios of cell-proliferation dye 670-labelled CAR Tregs. The CAR Tregs were pre-stimulated with HLA-A2-coated beads and IL-2. After co-culture for 72h, division of gated CPD-450^+^CD4^+^ or CPD-450^+^CD8^+^ responder T cells was determined by flow cytometry. **(A)** Schematic diagram of experiment setup. **(B)** Percent suppression was calculated using the ratio of the division index of the experimental condition to that in wells with no Tregs added. Representative **(B)** and averaged **(C)** data are shown. The NGFR and CD28wt Treg groups are repeated in the various graphs. n=3, averaged from two independent experiments. Statistics show overall effects two-way ANOVA with Holm-Sidak post-test comparing all constructs to CD28wt. **(D)** In vitro suppression of co-stimulatory molecule expression on dendritic cells. HLA-A2+ monocyte-derived dendritic cells were matured then co-cultured with the indicated type of CAR Treg (ratio of 1 dendritic cell to 2 CAR Tregs) for 72 hours. Flow cytometry analysis of mean fluorescence intensity of CD86 and CD80 on gated dendritic cells with representative and averages data shown. n=3-5, pooled from at least three independent experiments. Statistics show one-way ANOVA with Holm-Sidak post-test comparing all constructs to CD28wt. Mean ± SEM. **(E)** Correlation analyses comparing day 28 GVHD score to suppression of CD4 or CD8 T cell proliferation, or suppression of CD86 or CD80 co-stimulatory molecule expression on DCs. Suppression of CD86/CD80 expression was calculated using the ratio of MFI of the experimental condition over MFI of the control without Tregs (DC only). Statistics shown are Pearson correlation coefficient showing a line of best fit with 95% confidence intervals. * p < 0.05, ** p < 0.01, *** p < 0.001.

We then assessed the ability of the various CAR Tregs suppress dendritic cells (DCs) as determined by reduced expression of CD80 and CD86 (Hou et al., 2019; Wang et al., 2010). HLA-A2^+^CD14^+^ monocytes were cultured for 7 days in IL-4 and GM-CSF and matured for the last 2 days in a pro-inflammatory cytokine cocktail as described (Wang et al., 2010). Matured DCs were then co-cultured with CAR Tregs to test their ability to suppress expression of co-stimulatory ligands on DCs. Once again, we found that the most potent effect was mediated by the CD28wt-CAR, and that in this case the 4-1BB- and TNFR2-encoding CARs had a significantly lower ability to suppress expression of CD80 (**Figure 5D**). Expression of CD83 or HLA-DR were unaffected by CAR Tregs (data not shown).

In vitro suppression assays are often used to predict in vivo Treg function. We therefore sought correlations between our in vivo GVHD survival results (day 28 GVHD score), and our T-cell and/or DC suppression assay results (**Figure 5E**). While there was a trend towards a correlation between suppression of CD4^+^ and CD8^+^ T cell proliferation and CD86 expression on DCs and in vivo GVHD score, the only significant correlation was with suppression of CD80 on DCs.

### Gene expression analysis reveals CD28wt-CAR Treg are enriched in cell cycle and proliferation pathways compared to poorly functioning CAR Tregs

Finally, we asked how the various signaling domain CAR variants differed in their modulation of Treg gene expression. CAR Tregs were expanded, and purified as described above, before co-culture with A2-coated beads for 24h, then lysed for RNA sequencing. Controls consisted of ΔNGFR-transduced Tregs or Tconv either unstimulated or αCD3/28-stimulated with beads. Paired-end libraries were created from isolated RNA and sequences on a NextSeq 500 Illumina sequencer. Reads that mapped to the transgene sequences were filtered out to enable comparisons between groups and CAR-versus TCR-stimulation (**Supplemental Figure 7A**). Differential expression analysis was performed on the raw counts from filtered, aligned sequences. Within the Treg groups a number of differentially expressed transcripts were identified, with a heatmap depicting the top 50 differentially expressed genes shown in **Figure 6**. Strong patterns emerged, with clear similarities between unstimulated and PD-1-CAR-stimulated Tregs, as well as between CD28wt-CAR-, CD28mut-CAR- and TCR-stimulated Tregs.

**Figure 6.**
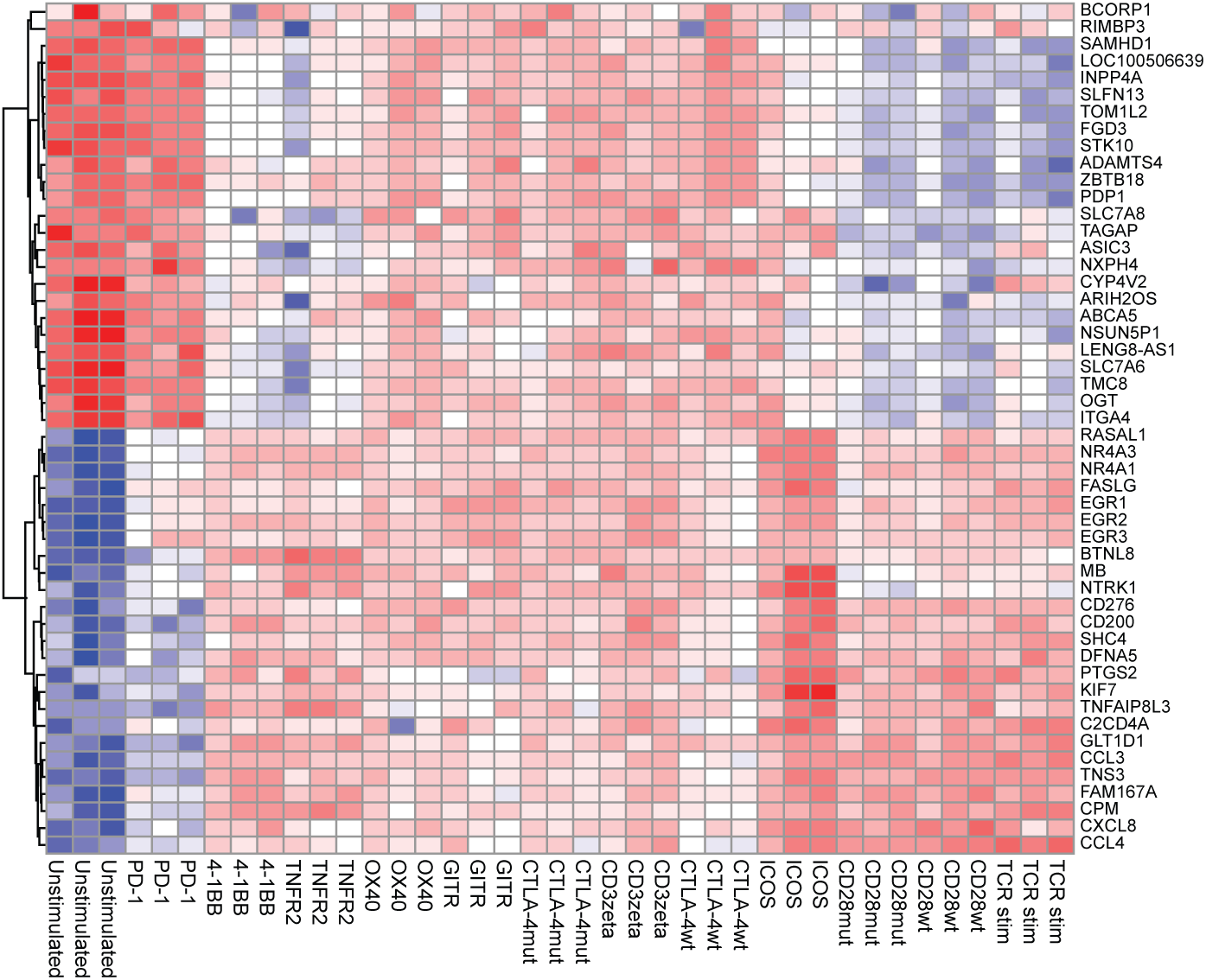
Effects of signaling domain CAR variants on the Treg transcriptome. Purified CAR Tregs were stimulated with HLA-A2-coated beads (CAR-stimulated) or anti-CD3/CD28 coated Dynabeads (TCR-stimulated) for 16 hours, then processed for RNA sequencing. Reads that mapped to transgene sequences were filtered out as described in Supplemental Figure 7A. Heatmap showing the top 50 differentially expressed genes compared with unstimulated Tregs.

Guided by the results from the GVHD study (**Figure 2**), we then asked if there were differences in the transcriptional profile between the CAR Tregs that performed the poorest - defined as groups where none of the mice survived to endpoints (CTLA-4wt and CTLA-4mut, PD-1, TNFR2, CD3zeta) and those performing best, bearing the optimal CD28wt-CAR (**Figure 7A, Supplemental Table 1**). We found that CD28wt-CAR Treg gene expression was highly enriched in several genes targeted by transcription factors associated with cell proliferation and DNA replication, such as the MCM complex, MYB and E2F4. NFKB1 and RELA transcription factor targets were also elevated, indicating that the NFκB signaling pathway may be important for cell cycling and proliferation secondary to CAR activation. Pathway analysis showed that CD28wt-CAR Tregs also expressed elevated levels of transcripts associated with mTORC1 signaling, G2M checkpoint and Myc targets, supporting the concept that cell cycling is an important component of CAR-Treg function.

**Figure 7.**
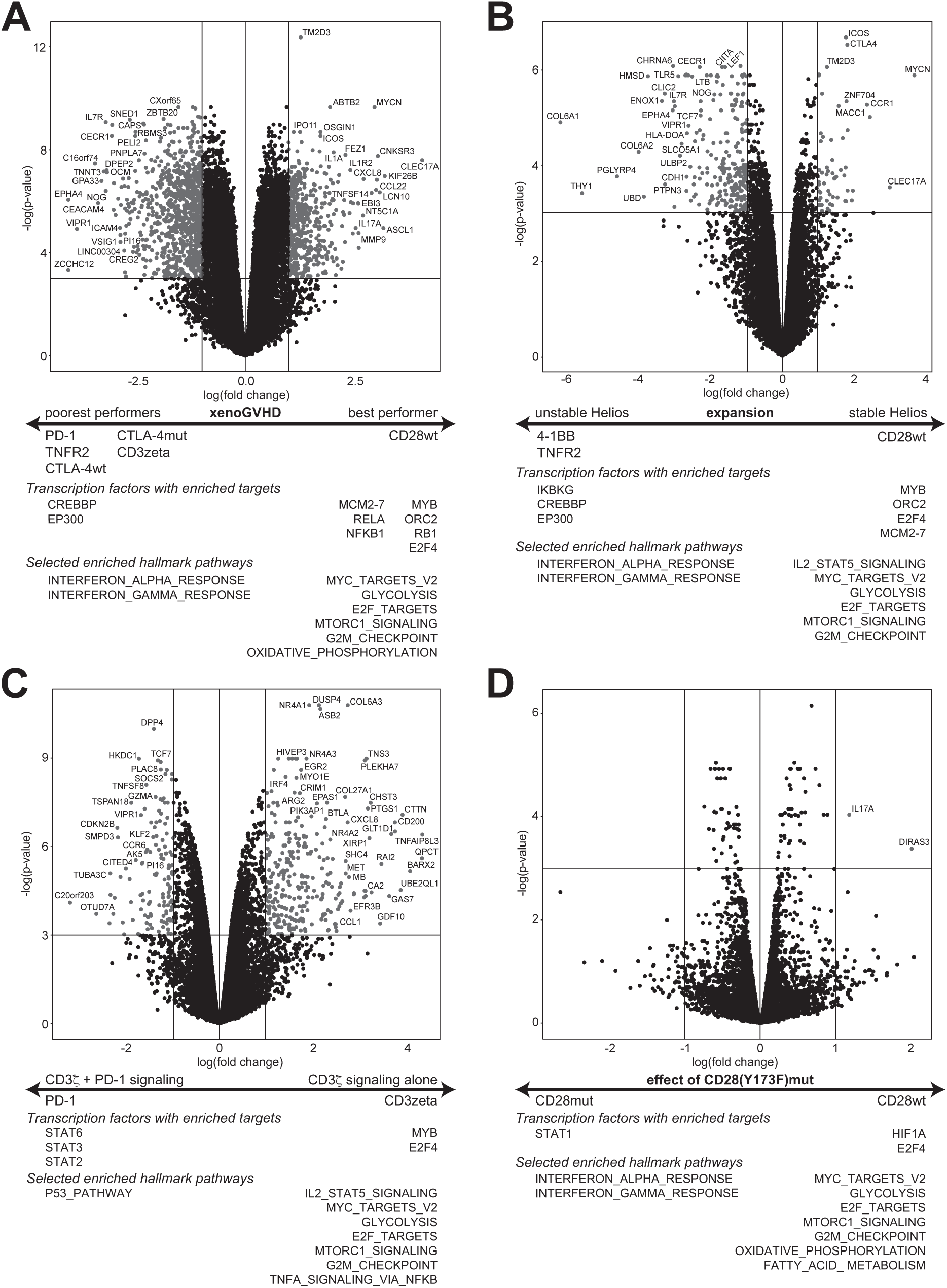
Effects of signaling domain CAR variants on the Treg transcriptome. CAR Tregs were prepared and stimulated as in Figure 8. **(A)** Differentially expressed transcripts between CD28wt and PD-1, TNFR2, CTLA-4wt, CTLA-4mut, CD3zeta, highlighting differences in the transcriptome between CD28wt-CAR Tregs and the CAR Treg groups for which no mice remained at the experimental endpoint (day 42) of the GVHD study. **(B)** Differentially expressed transcripts between CD28wt-CAR Tregs and 4-1BB/TNFR2-CAR Tregs to highlight differences between constructs that stimulated proliferation upon CAR-stimulation and either maintained or destabilized Treg phenotype. **(C)** Differentially expressed transcripts between CD3zeta-(first-generation CAR) and PD1-CAR Tregs, highlighting the negative effects of PD-1 signaling on CD3zeta signaling in Tregs. **(D)** Differentially expressed transcripts between CD28wt- and CD28mut-CAR Tregs highlighting differences caused when stimulated via mutated CD28(Y173F) in Tregs. Volcano plots (above) and selected results from enrichment analysis of transcription factor targets and hallmark pathways (below). Full lists of differentially expressed transcription factor targets and hallmark pathways with normalized enrichment scores and adjusted p values are found in **Supplemental Tables 1-4**. n=3 from two independent experiments.

CD28wt-, 4-1BB- and TNFR2-CAR Tregs all proliferated in response to A2, but only CD28wt-CAR Tregs maintained Helios expression (**Figure 5**), so we queried how the transcriptional profiles of 4-1BB- and TNFR2-CAR Tregs differed from that of the CD28wt-CAR Tregs (**Figure 7B, Supplemental Table 2**). We found that in comparison to the 4-1BB- and TNFR2-CARs, the CD28wt-CAR had higher expression of ICOS and CTLA-4, but lower IL7R (CD127). As in **Figure 7A**, pathway analysis revealed that the CD28wt-CAR Tregs expressed elevated levels of genes of transcription factor targets and pathways associated with cell cycle and division. On the other hand, Tregs bearing 4-1BB- and TNFR2-CARs expressed elevated levels of genes in inflammatory cytokine response pathways (IFN and TNF) and of CREBBP and EP300 transcription factor targets. Targets of IKBKG, a transcription factor associated with suppression of the NFκB pathway, were also elevated in 4-1BB and TNFR2 samples.

Interestingly, we noticed that the top differentially expressed genes in the PD-1-CAR Tregs were more similar to those of unstimulated cells than the CD3zeta-CAR, a first-generation CAR construct (**Figure 7C**). We therefore performed an enrichment analysis to determine the pathways stimulated by PD-1 that override CD3zeta activating signals (**Figure 7C, Supplemental Table 3**). In accordance with an activated phenotype, CD3zeta had higher expression of transcripts associated with cell cycling and proliferation. We found that the PD-1-CAR was enriched for genes associated with the p53 tumor suppressor gene pathway, providing a potential mechanism underlying inhibitory PD-1 signaling function in Tregs.

Since the CD28mut-CAR Tregs failed to proliferate after CAR stimulation in vitro, and, in comparison to CD28wt-CAR Tregs, had a diminished function in vivo, we next interrogated the transcriptional differences between these two cell types (**Figure 7D, Supplemental Table 4**). Our data showed that there were only two differentially expressed genes, *IL17A* and *DIRAS3*, but that CD28wt-CAR Tregs gene expression was relatively enriched in pathways related to cell cycle and metabolism. These data indicate that the functional differences between these CARs is likely related to differences in post-transcriptional events (e.g. protein phosphorylation) which are not evident from short term gene expression analysis.

### Differences in CAR- versus TCR-stimulated gene expression in Tregs and Tconv cells

To date there has been no analysis of how CAR-stimulated gene expression compares to TCR stimulation, in either Tregs or Tconv cells. We thus also compared the transcriptional profiles of CD28wt-CAR and αCD3/28-stimulated Tregs (**Supplemental Figure 7B, Supplemental Table 5**), finding an overall similar pattern of gene expression, with only a handful of differentially expressed genes. Gene set enrichment analyses revealed that both stimuli upregulated transcripts associated with proliferation, but gene expression of CD28wt-CAR stimulated Tregs was enriched in genes associated with response to pro-inflammatory cytokines. These data indicate that, at least for the anti-A2 scFv, stimulation via the CD28wt-CAR in Tregs faithfully replicates endogenous TCR/CD28 signaling activated by αCD3/28 crosslinking. We repeated this comparison using CAR Tconv cells and noted an enrichment of pro-inflammatory cytokine gene expression, e.g. *IL2, IL13*, and *IL5*, and genes associated with cytokine response in CAR-stimulated versus TCR-stimulated cells (**Supplemental Figure 7C, Supplemental Table 6**). Conversely, expression in TCR-stimulated cells was enriched in genes associated with proliferation and DNA replication. Interestingly, like Tregs, CAR-stimulation in Tconvs upregulated transcripts associated with NF-κB-related transcription factors, e.g. NFKB1, REL and RELA, which could be driven by differences in the composition and structure of the immunological synapse between CAR- and TCR-ligation (Davenport et al., 2018).

We also compared gene expression stimulated by the CD28wt-CAR expressed in Treg and Tconvs (**Supplemental Figure 7D, Supplemental Table 7**), revealing a large number of differentially expressed genes, many of which were consistent with the expected Treg and Tconv phenotypes. For example, The CD28wt-CAR Tregs were enriched in expression of *FOXP3*, *IKZF2* (Helios) and numerous genes associated with suppression, such as *CTLA4*, *FCRL3*, *LRRC32* (GARP), and *TIGIT* (Bin Dhuban et al., 2015; Josefowicz et al., 2012). CAR Tregs also exhibited upregulated transcription of genes associated with TGFβ, IL2/STAT5, and NF-κB signaling. Tconv expressed a number of effector cytokines, e.g. *IL2*, *IL5*, *IFNG*, *IL13*, and *IL9*. Interestingly, this comparison showed that gene expression in CD28wt-CAR Tconvs was enriched in many of the same proliferation-related transcription factor targets that were enriched in the CD28wt-CAR Tregs when compared to poorly functional CAR Tregs. These data suggest that although proliferation is an important feature of functional CAR Tregs, it is less prominent than in CAR Tconvs.

## Discussion

Here we report the first comprehensive comparison of co-receptor signaling domain CAR variants in Tregs, revealing key features of CAR Treg biology and discovering in vitro assays which do, or do not, correlate with in vivo function. Surprisingly, and distinct from results with CAR Tconvs, we found that inclusion of the wild type CD28 co-stimulatory domain was essential for potent function. No other CD28-superfamily member tested could substitute for CD28; even a version of CD28 with a single point mutation was inferior. Moreover, CARs encoding domains from TNFR family members were unable to confer a significant protective effect in comparison to irrelevant-antigen-specific control Tregs. These data show that Tregs and Tconv cells have distinct requirements for optimal CAR-mediated suppression/effector function and reveal new aspects of Treg biology that can be used to further optimize CAR design.

Importantly, no single in vitro assay was absolutely predictive of the in vivo effect of CAR Tregs. Of particular note was the limited correlation between in vivo function and three in vitro assays which are commonly used to gauge human Treg function: CAR-mediated stimulation of activation/functional markers, proliferation, and in vitro suppression of T cells. Rather, the in vitro assay that most strongly correlated with GVHD score at day 28 was suppression of CD80 expression on mature DCs. Notably, CD80 has a higher affinity for CTLA-4 than does CD86 (Walker and Sansom, 2011) and CAR-stimulated CD28wt-CAR Tregs had high and prolonged CTLA-4 expression. Taken together, these data support the notion that APCs play a key role in mediating and propagating Treg suppression and suggest a primary mechanism of CAR Treg function is via CTLA-4-mediated suppression. Our data also indicate that strong CAR-stimulated proliferation is essential for in vivo function, providing that lineage stability is not compromised.

It has been shown previously that the level of CAR expression affects function (Guedan et al., 2018), leading to the possibility that some of our findings could be related to the observed differences in mean fluorescence intensity of CAR expression. Indeed, all but the CD28mut- CAR had significantly lower MFIs than the CD28wt CAR, as well as diminished abilities to stimulate expression of key Treg effector proteins (*i.e*. LAP, GARP and CTLA-4). It seems unlikely, however, that differences in expression could solely account for the observed differences, since many CARs were capable of stimulating in vitro suppression of T cell proliferation and dendritic cell expression of CD80/86 and, despite being expressed at the same level as the CD28wt-CAR, the CD28mut-CAR Tregs were unable to proliferate in response to A2-mediated stimulation in vitro. Moreover, although the ICOS- and OX40-CARs were expressed at significantly lower levels than the CD28wt-CAR, at high ratios they both provided a survival benefit in vivo, in comparison with mice treated with HER2-CAR Tregs.

We were surprised to find such a clear difference between the CD28wt-CAR and all other CAR constructs tested in the GVHD model, an effect that was particularly striking at low Treg:PBMC ratios. Although all of the in vitro assays showed trends towards the superiority of the CD28wt-CAR, no single in vitro assay could have predicted this in vivo outcome. The CD28wt-CAR Tregs clearly survived better in vivo and indeed, at day 7, this group had a significantly higher absolute number of CAR^+^FOXP3^+^Helios^+^ cells in circulation than any other groups. The observed survival advantage of the CD28wt-CAR Tregs in vivo is supported by the transcriptome analysis which showed an enrichment for many genes in cell cycle/proliferation/DNA replication related pathways (e.g. ORC, MYC and the MCM complex), as well as by the ability to proliferate in response to A2 in vitro. However, the mere ability to stimulate proliferation is clearly not sufficient for full function, since the proliferative signals activated by the 4-1BB and TNFR2-CARs led to loss of TSDR methylation and Helios expression.

Notably, the distinction between the downstream effects of CD28wt- and 4-1BB/TNFR2-CAR-stimulated proliferation was evident in terms of differential stability of Helios, but not of FOXP3, expression. In mice, Helios is dispensable for Treg development but is known to be required for stability and suppressive function. For example Helios-deficient Tregs are unable to control peripheral immune effector cells or prevent colitis (Kim et al., 2015a; Nakagawa et al., 2016; Skadow et al., 2019; Thornton et al., 2019). The rapid loss of Treg stability in 4-1BB- or TNFR2-CARs could possibly be explained by preferential expansion of contaminating non-Treg cells, but this seems unlikely given that TSDR analysis prior to CAR stimulation showed all cell populations had an almost completely demethylated TSDR.

Interestingly, transcriptome analysis revealed a strong enrichment in expression of the NF-κB pathway/target genes in CD28wt-CAR Tregs. This finding is consistent with the fact that CD28 co-stimulation activates the NF-κB pathway via Protein kinase C (Cheng et al., 2011; Takeda et al., 2008) and that both p65 (Rela) and c-Rel (Rel) have important roles in Treg development, maintenance, and function (Grinberg-Bleyer et al., 2017; Long et al., 2009; Messina et al., 2016; Oh et al., 2017). We also detected a decrease in the level of CREBBP (CBP) and EP300 (p300) transcriptional targets. The CBP/p300 complex contains a bromodomain that binds acetylated histones on the *FOXP3* promoter, a process shown to be critical for promotion of Treg development and stability (Du et al., 2013; Ghosh et al., 2016; Nagai et al., 2019), However, we did not detect FOXP3 destabilization or defects in other Treg functional properties, so the observed changes in gene expression related to this pathway were not sufficient to result in a functional Treg defect.

Given the extensive evidence for the benefit of 4-1BB encoding CARs in anti-tumour-CAR T cell function, we were surprised to find that inclusion of this domain provided no apparent benefit for Tregs. This finding is consistent with a recent report from Boroughs et al who used a CD19 CAR system to compare the function of first generation, CD28 or 4-1BB encoding CARs in Tregs (Boroughs et al., 2019). They found that a 4-1BB CD19-CAR stimulated expression of CTLA-4 and LAP but had diminished suppressive function in vitro. Previous reports on the function of 4-1BB in Tregs have been conflicting with some evidence showing promotion of Treg proliferation and suppression (Lee et al., 2005; Sun et al., 2002; Zhang et al., 2007), and others showing that 4-1BB signaling inhibits Treg suppression (Akhmetzyanova et al., 2016; Buchan et al., 2018; Smith et al., 2011).

We were also surprised to find no benefit for inclusion of a TNFR2 domain, despite extensive evidence for the beneficial effect of TNFR2 ligation on Treg proliferation in other systems (He et al., 2016; Lam et al., 2017; Yang et al., 2018). Although both 4-1BB- and TNFR2-CAR Tregs proliferated in response to A2 in vitro, they did not have a survival advantage in vivo. Relative to the CD28wt-CAR, 4-1BB- and TNFR2-CAR Tregs upregulated transcriptional pathways involved in type I and type II interferon response and TNFα signaling, and down regulated those associated with IL2/STAT5 signaling, glycolysis and mTORC1 signaling, findings consistent with the observed destabilized Treg phenotype. Moreover, 4-1BB and TNFR2 had higher relative expression of targets of IKBKG, which is involved in the negative regulation of NF-κB signaling, further suggesting that this axis is important in CAR Treg function.

Both GITR and OX40 have well-established roles in Treg development in the thymus (Mahmud et al., 2014), but their roles in mature Tregs are unclear and nuanced. Some reports show that ligation of OX40 or GITR decreases Treg suppressive capability (Cuzzocrea et al., 2005; Kitamura et al., 2009; Shimizu et al., 2002; Vu et al., 2007; Xiao et al., 2012; Zhang et al., 2018) with others reporting a benefit for Treg proliferation and function (Ephrem et al., 2013; Kinnear et al., 2013), possibly related to the kinetics of receptor stimulation. We found that although the GITR- and OX40-CAR Tregs provided an intermediate GVHD survival benefit at the high Treg:PBMC ratio in vivo, neither CAR was able to stimulate proliferation or strong activation of Tregs in vitro. Poor proliferation with the OX40-CAR is consistent with the finding that stimulation of Treg expansion via anti-CD3 and anti-OX40 antibodies is inferior to treatment with anti-CD3 and anti-CD28 antibodies (Golovina et al., 2008).

Focusing on CARs encoding inhibitory co-receptors, inclusion of PD-1 completely blocked CD3ζ-mediated activation of Tregs, with no A2-stimulated expression of activation markers or effector molecules, proliferation or suppression. Transcriptome analysis comparing the PD-1-CAR to a first generation CD3ζ-CAR revealed that PD-1 signaling in Tregs resulted in upregulated p53 signaling, a pathway strongly associated with cell cycle arrest (Kastenhuber and Lowe, 2017). While a strong association between PD-1 signaling and development of peripherally-induced Tregs has been established (Chen et al., 2014), whether there is a functional role for PD-1 signaling in thymic-derived Tregs, and, by extension, CAR Tregs, is less clear. However, a recent report showed that PD-1 blockade augments proliferation and suppressive activity of Tregs, which is consistent with our activation data showing that stimulation via PD-1 limited CAR Treg ability to activate, proliferate and suppress T cell proliferation (Kamada et al., 2019). In terms of CTLA-4, although clearly required for Treg function (Walker and Sansom, 2011), the extent of its requirement for cell-intrinsic signaling, as against cell-extrinsic effects is unclear (Walker and Sansom, 2015). Our data confirm that the point mutation Y165G improves cell surface expression (Nakaseko et al., 1999) and show that CTLA-4 intrinsic signaling is not beneficial for Tregs.

A notable limitation of our study is due to the paucity of models in which to test human Treg function in vivo. We elected to use the well-established GVHD model because it is a system in which the advantage of CAR Tregs has previously been observed (Dawson et al., 2019; MacDonald et al., 2016) and it is significantly more high throughput than other humanized mouse models, and thus amenable to testing multiple CAR Treg groups in parallel. Mechanistically, it is known that the suppressive effects of Tregs in this model are at least partially due to TGF-β (Lienart et al., 2018) and CTLA-4 signaling (Zaitsu et al., 2017), two pathways that are strongly stimulated by the CD28wt-CAR. However, we acknowledge that this model does not recapitulate a normal immune response, since there is poor engraftment of human APCs, the lack of a lymph node network, and only selective cross-reactivity of mouse with human homing stimuli/receptors to support human T cell trafficking. In the context of a complete immune system, different CAR-stimulated effects may be observed, a possibility that will require study in systems without the limitations of the GVHD model.

Collectively, our findings support the use of CD28wt-based CAR for use in Treg therapies. Future studies could seek to optimize CAR signaling moieties by using third-generation CARs incorporating CD28 or adding additional signals that reinforce Treg identity. Our extensive analysis of the in vitro and in vivo properties and effects of signaling-domain CAR variants provides a platform from which to design future studies and also leads to significant insight into the essential properties of engineered Treg therapies.

## Acknowledgments

This work was supported by grants from the Canadian Institutes of Health Research (CIHR) FDN-154304 and TxCell (to MKL). NAJD is supported by a CIHR doctoral award, IRS is supported by a salary award from the UBC School of Biomedical Engineering, VCWF received an American Society for Transplantation Student Internship Research Program salary award and MKL receives a salary award from the BC Children’s Hospital Research Institute.

## Author contributions

NAJD conceived, designed and conducted experiments, analyzed data, and wrote the manuscript. IRS designed and conducted experiments, analyzed data, helped write and critically review the manuscript. GN analyzed the RNA sequencing data. VF, QH, EM, GS JG, and MS conducted experiments and analyzed data. PCO provided intellectual input, and critically reviewed the manuscript. MM provided intellectual input and logistical support and critically reviewed the manuscript. MKL secured funding, conceived and designed experiments, provided overall direction, and wrote the manuscript.

## Declaration of Interests

The authors of this manuscript have received research funding from TxCell SA to partially support this work.

## Material and Methods

### Generation of signaling domain CAR variants

Sequences for the intracellular region of selected co-stimulatory or co-inhibitory molecules were scraped from Uniprot (Consortium, 2018), then codon optimized for *Homo sapiens* using the codon optimizer from Thermo Fisher Scientific Invitrogen GeneArt Gene Synthesis service. gBlocks® Gene Fragments were ordered from Integrated DNA Technologies (Coralville, Iowa) with appropriate unique restriction enzyme recognition sequences to clone in frame with an anti-HER2 or anti-HLA-A2 scFv (derived from the BB7.2 mAb) (MacDonald et al., 2016) in place of the existing CD28wt co-stimulatory domain. From N- to C-terminal, the CAR construct contained: the scFv, a myc-tag, a stalk region from human CD8α, the specified human transmembrane and intracellular co-receptor domain sequences, then CD3ζ. In some cases we also used a first-generation CAR construct which encoded CD3ζ directly C-terminal of the CD28 transmembrane region. All CAR constructs were cloned into a lentivirus backbone containing a bicistronic promoter with the minimal-CMV promoter controlling expression of the truncated nerve-growth-factor receptor (ΔNGFR) as a transduction marker and the EF1α promoter controlling expression of the CAR construct. Surface expression was determined by flow cytometry with transiently transfected HEK 293T cells (jetPRIME®, Polyplus Transfection). Viral particles were produced as described (Allan et al., 2008).

### Treg sorting, transduction, and expansion

CD4^+^ T cells were isolated from PBMCs (HLA-A2-negative donors) using RosetteSep (STEMCELL Technologies, 15062), then enriched for CD25^+^ cells (Miltenyi, 130-092-983) prior to sorting live CD4^+^CD25^hi^CD127^lo^ Tregs (for most experiments) or CD4^+^CD127^lo^CD25^hi^ CD45RA^+^Tregs (for long-term in vitro culture experiments) using a MoFlo® Astrios (Beckman Coulter) or FACSAria II (BD Biosciences). In experiments where conventional T cells (Tconv) were used, the live CD4^+^CD25^lo^CD127^hi^ fraction was also sorted. Sorted cells were stimulated with L cells and αCD3 mAb (OKT3, UBC AbLab; 100ng/mL) in Immunocult-XF T cell media (STEMCELL Technologies, 10981) with 1000U/ml of IL-2 (Proleukin) as described (MacDonald et al., 2016). One day later, cells were transduced with lentivirus at a multiplicity of infection of 10 virus particles:1 cell. At day 7, ΔNGFR^+^ cells were purified with magnetic selection (Miltenyi, 130-091-330) then used in assays. For in vivo experiments cells were re-stimulated with L cells as above and expanded for an additional 5 days prior to injection.

### Flow cytometry

For phenotypic analysis, cells were stained with fixable viability dye (FVD, Thermo Fisher Scientific, 65-0865-14; Biolegend, 423102) and for surface markers before fixation and permeabilization using eBioscience FOXP3/Transcription Factor Staining Buffer Set (Thermo Fisher Scientific, 00-5523-00) and staining for intracellular proteins. Samples were read on a Cytoflex (Beckman Coulter) and results analyzed using FlowJo Software version 10.5.3 (Tree Star).

Surface staining was performed for NGFR (Miltenyi, 130-091-885; BD Biosciences, 562122, 562562), myc (9E10 clone, UBC Ablab, Vancouver, Canada), CD4 (Biolegend, 317410, 300558), CD25 (Miltenyi, 130-091-024), LAP (Thermo Fisher Scientific, 25-9829-42), GARP (BD Biosciences, 563958), CD69 (Biolegend, 310945, 310931), CD71 (BD Biosciences, 563768) and CD127 (Thermo Fisher Scientific, 48-1278-42), CD39 (Biolegend, 328214), CD80 (BD Biosciences, 557226), CD11c (BD Biosciences, 555392), CD86 (BD Biosciences, 555660), CD83 (Biolegend, 305324), HLA-DR (Biolegend, 307646). Intracellular staining was performed for CTLA-4 (BD Biosciences, 563931), FOXP3 (Thermo Fisher Scientific, 25-4777-42, 12-4777-42), and Helios (Biolegend, 137223) using the eBioscience FOXP3/Transcription Factor Staining Buffer Set (Thermo Fisher Scientific, 00-5523-00).

For in vivo experiments, 50μL of blood was collected weekly and at endpoints. Ammonium chloride was used for red blood cell lysis. Cells were resuspended in PBS with anti-mouse CD16/32 (Thermo Fisher Scientific, 14-0161-82) and stained for extracellular markers using fixable viability dye (FVD; Thermo Fisher Scientific, 65-0865-14), anti-mouse CD45 (Thermo Fisher Scientific, 56-0451-80), and anti-human CD45 (BD Biosciences, 560777), CD4 (Biolegend, 300554), CD8 (Thermo Fisher Scientific, 48-0087-42), anti-human CD271 (NGFR; BD Biosciences, 562562), HLA-A2 (Biolegend, 343306), myc (9E10 clone, UBC Ablab, Vancouver, Canada). Intracellular staining for FOXP3 (Thermo Fisher Scientific, 25-4777-42) and Helios (Biolegend, 137223) was done with the eBioscience FOXP3/Transcription Factor Staining Buffer Set (Thermo Fisher Scientific, 00-5523-00). 10,000 counting beads were added to every sample (Thermo Fisher Scientific, 01-1234-42). The gating strategies for the xenogeneic GvHD experiments are illustrated in **Supplemental Figure 2B.**

### Xenogeneic graft-versus-host disease

8- to 12-week-old female NSG mice (The Jackson Laboratory, Maine USA; bred in house) received whole-body irradiation (150 cGy, RS-2000 Pro Biological System) 1 day before injection of 10 × 10^6^ HLA-A2^+^ PBMCs with or without 5 × 10^6^ (high ratio) or 2.5 × 10^6^ (low ratio) CAR Tregs intravenously into the tail vein. Saline-injected (PBS) mice served as a control and PBMC-only and CD28wt CAR conditions were included in every experimental cohort. GVHD was scored based on weight, fur texture, posture, activity level, and skin integrity, with 0 to 2 points per category as described (Cooke et al., 1996; Hill et al., 1997). Peripheral blood from the saphenous vein was centrifuged; then erythrocytes were lysed and leukocytes were measured by flow cytometry.

### In vitro functional assays

To test the effects of CAR or TCR-mediated stimulation, Tregs were cultured with limiting IL-2 (100U/mL) for 24 hours, then re-counted and co-cultured with irradiated anti-CD3/anti-CD28-loaded CD64-expressing K562 cells, HLA-A*24:02-expressing K562 cells, HLA-A*02:01-expressing K562 cells (Dawson et al., 2019) or HLA-A*02:01 FlowPRA Single Antigen beads (One Lambda, Thermo Fisher Scientific) at the specified ratios and timepoints. The determine amounts of cytokine profile, supernatants from stimulated CAR Tregs were taken diluted 2-fold before analysis via cytometric bead array (BD Biosciences, 560484) using the manufacturer’s recommended protocol.

### Suppression of T cells

Allogeneic HLA-A2^−^ CD3+ responder cells were labeled with cell proliferation dye eF450 (Thermo Fisher Scientific, 65-0842-85), then stimulated with anti-CD3/CD28 Dynabeads (Thermo Fisher Scientific, 11141D) at a ratio of 1:2 (bead to cell) in X-Vivo 15 medium (Lonza, 04-744Q) supplemented with 5% human serum (Wisent Bio Products, 022210), 1% glutamax (Gibco, 35050-061), 1% penicillin/streptomycin (Gibco, 15140-122), hereafter referred to as “XH medium”, with 100U/mL IL-2 (Proleukin). Simultaneously, 2 × 10^5^ CAR Tregs, or ΔNGFR (no CAR) Treg control cells, were labeled with cell proliferation dye eF670 (Thermo Fisher Scientific, 65-0840-85), and stimulated with a 1:1 ratio of HLA-A*02:01 FlowPRA Single Antigen beads (One Lambda, Thermo Fisher Scientific) in XH medium with 100U/mL IL-2. After 24-hour incubation, half of the volume of Tregs + beads was serially diluted in XH medium with 100U/mL IL-2. Dynabeads were removed from CD3/28-stimulated CD3+ responder cells by resuspension and 3-5-minute incubation on a magnet (STEMCELL Technologies, 18103). Bead-free responder cells were then resuspended in fresh XH media + 100U/mL IL-2 and 0.5 × 10^5^ cells were plated on top of each well. The co-cultures were incubated for 3 additional days, then analyzed by flow cytometry. Percent suppression was calculated based on the division index of gated CD4+ or CD8+ responder T cells in comparison to the division index in the absence of Tregs as described (McMurchy and Levings, 2012). Percent suppression was calculated as follows: 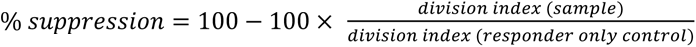.

### Suppression of antigen-presenting cells

CD14+ monocytes were isolated (STEMCELL Technologies, 18058) from HLA-A2+ human PBMC, then cultured in X-Vivo15 medium supplemented with 5% human serum (Wisent Bio Products, 022210), 1% glutamax (Gibco, 35050-061), 1% penicillin/streptomycin (Gibco, 15140-122), 1mM sodium pyruvate (STEMCELL Technologies, 07000), herein referred to as “DC medium”, and 100ng/mL IL-4 (STEMCELL Technologies, 78045.1), 50ng/mL GM-CSF (STEMCELL Technologies, 78015) for 5 days (media and cytokines refreshed on day 3). Cells were matured using DC medium containing a cytokine cocktail of 100ng/mL IL-4, 50ng/mL GM-CSF, 10ng/mL TNF-α (Thermo Fisher Scientific, 14-8329-63), 1μg/mL PGE-2 (Sigma, P6532), 10ng/mL IL-1β (STEMCELL Technologies, 780341) and 100ng/mL IL-6 (Thermo Fisher Scientific, 14-8069-80) for 2 additional days. In the last 24 hours of culture, 1000U/mL IFN-γ (Thermo Fisher Scientific, 34-8319-82) was added. Matured cells were resuspended in DC medium with a final concentration of 50U/mL IL-2 and co-cultured with CAR Tregs at a ratio of 1 dendritic cell to 5 CAR Tregs for 4 days then analyzed by flow cytometry.

### Generation and analysis of stimulated CAR Tregs via RNA sequencing

NGFR-purified CAR Tregs and CAR Tconv cells (2.5 × 10^5^) were stimulated using 1.25 × 10^5^ HLA-A*02:01 FlowPRA Single Antigen beads or 2.5 × 10^5^ anti-CD3/CD28 Dynabeads for 24 hours in Immunocult-XF T cell media (STEMCELL Technologies, 10981). Approximately 0.5 × 10^5^ cells were used for flow cytometry analysis and total RNA was isolated from the remaining cells using the protocol to omit small RNAs (New England Biolabs, T2010S). RNA was quantified using fluorometer (Thermo Fisher Scientific, Q32855) and quality (RNA Integrity Number) determined using Bioanalyzer analysis (Agilent, 5067-1513). All RNAs had an RNA integrity number > 8.0. mRNA enrichment and library preparations were performed with a NEBNext Poly(A) mRNA Magnetic Isolation Module and a NEBNext Ultra II Directional RNA Library Prep Kit for Illumina (both New England Biolabs). Paired-end libraries were sequenced (43 × 43 bp reads) on a NextSeq 500 (Illumina).

Reads that mapped to the transgene using the data processing method outlined in (**Supplemental Figure 7A**) were removed. The read sequences were aligned to both the GRCh37/hg19 reference genome using STAR (v2.5.0a) and the transgene sequences using Burrows-Wheeler Aligner (v0.7.10). Cross-referencing the alignment files (.bam) from both reference genome and transgene sequences allowed for reads that mapped to the transgene sequences to be filtered out from further analysis by using unique identifiers for each read. In R, raw count matrices were generated using HTSeq (v0.11.2), then scale factors were calculated to take into account differences in library sizes using edgeR (v3.24.3) and normalization was performed using limma (v3.38.3) as in (Law et al., 2016). Log(CPM) and visualization was performed using: ggplot2 (3.1.1), RColorBrewer (v1.1.2), tibble (2.1.1), pheatmap (v1.0.12), stats (v3.5.1), and gplots (v3.0.1.1). The code used for data analysis is available on GitHub: https://github.com/najdawson/sigDom-RNAseq-git/

### Statistical analysis

All statistics were done using Prism 8.1.1. For all studies, normality was assumed. Corrections for multiple comparisons were made as described in each figure.

### Study Approval

For all studies, healthy volunteers gave written informed consent according to protocols approved by the University of British Columbia Clinical Research Ethics Board (UBC-CREB) and Canadian Blood Services. Commercial leukapheresis blood products were purchased from STEMCELL Technologies (Vancouver, Canada). Animal protocols were approved by the UBC Animal Care Committee. Some schematic diagrams were created with Biorender (http://biorender.com) and used with permission.

