## Supplemental Figures & Tables for "Functional effects of chimeric antigen receptor co-receptor signaling domains in human Tregs"

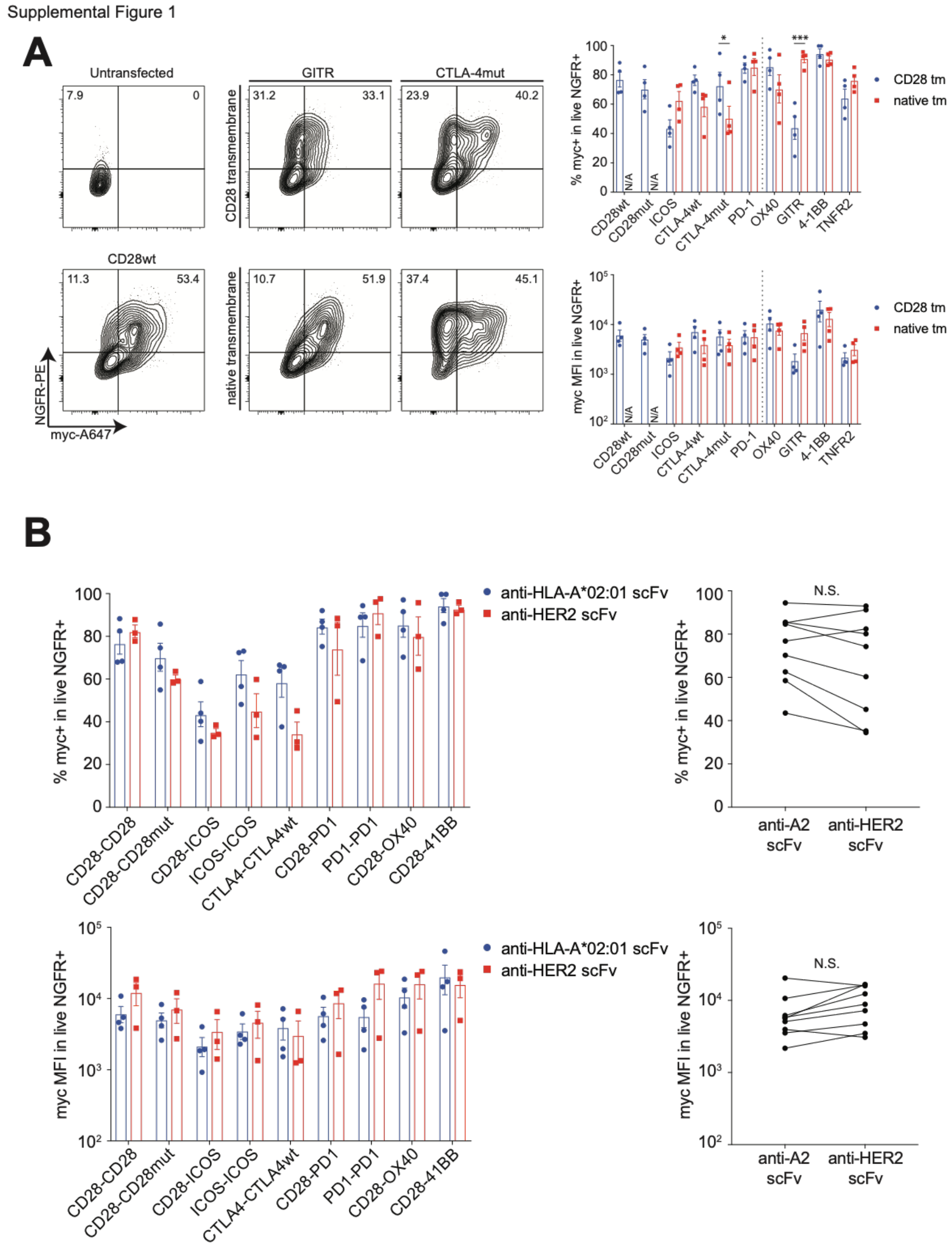


**Supplemental Figure 1. Surface expression of signaling domain CAR variants with different transmembrane domains or scFvs. (A)** Anti-HLA-A2 CAR variants were generated differing in their co-stimulatory domains, with either their original (or "native") transmembrane domains or a CD28 transmembrane domain. These constructs were transiently transfected into 293T cells and assessed by flow cytometry 48 hours after transfection, for surface expression of the CAR, evidenced by myc-tag, and the transduction marker, truncated NGFR (CD271). Left: representative flow cytometry plots. Right: summarized data of percent of mean fluorescence intensity of myc-tag detection in the live NGFR+ fraction. Data are n=3-4, pooled from at least three independent experiments. **(B)** scFvs from selected anti-HLA-A2 CAR constructs from (A) were replaced with an anti-HER2 scFv, and resultant constructs were transiently transfected into 293T cells and assessed by flow cytometry 48 hours after transfection for surface expression of the CAR, evidenced by myc-tag, and the transduction marker, truncated NGFR (CD271). Top: percent myc positive cells in live NGFR+ cells. Bottom: mean fluorescence intensity of myc in live NGFR$^+$ cells. Individual data (left) and summarized data (right) are shown. Statistics show one-way ANOVA with Holm-Sidak post-test comparing each transmembrane variant or scFv pair. Mean ± SEM. * $p < 0.05$, *** $p < 0.001$. "n.s." denotes not significant.

Supplemental Figure 2

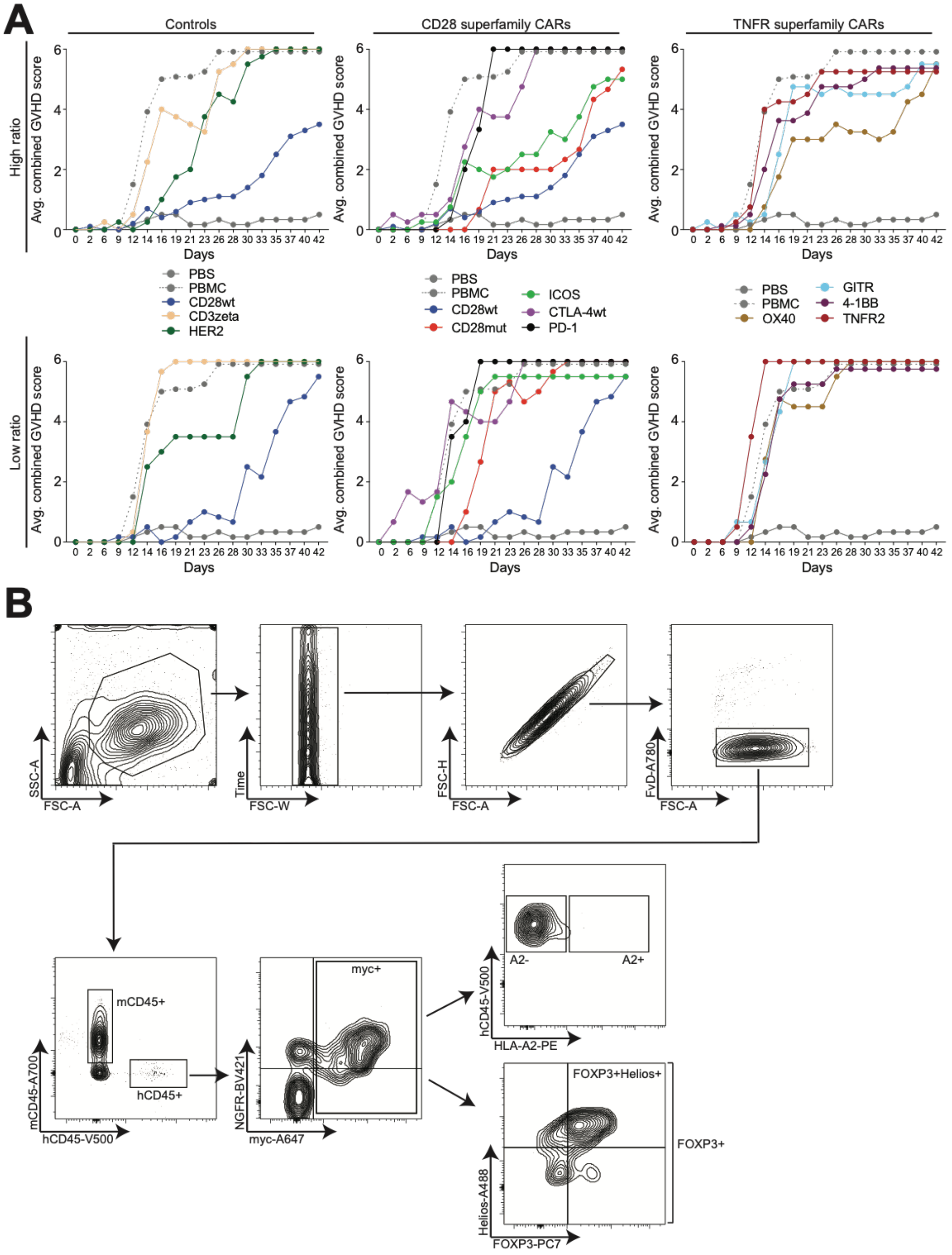


**Supplemental Figure 2. Supporting information for GVHD model. (A)** GVHD score for mice in GVHD model. Mice were scored on a scale from 0-3 for the following factors: weight loss, skin inflammation, fur maintenance, pain, deteriorating posture. If a score of 3 in any category or a combined score of 6 occurred, the mouse was sacrificed. Plotted are the average rolling combined GVHD scores for each experimental group, split by CAR signaling-domain superfamily and Treg:PBMC ratio. If a mouse was sacrificed for a score of 3 in one category, a value of 6 was assigned. Numbers of replicates are as in Figure 2, pooled from three independent experiments. Means without error are shown. **(B)** Gating strategy for flow cytometry analysis of GVHD experiments. Gates are as marked. Gating of CD28wt CAR Tregs from a high ratio mouse at day 7 is shown as an example.

Supplemental Figure 3

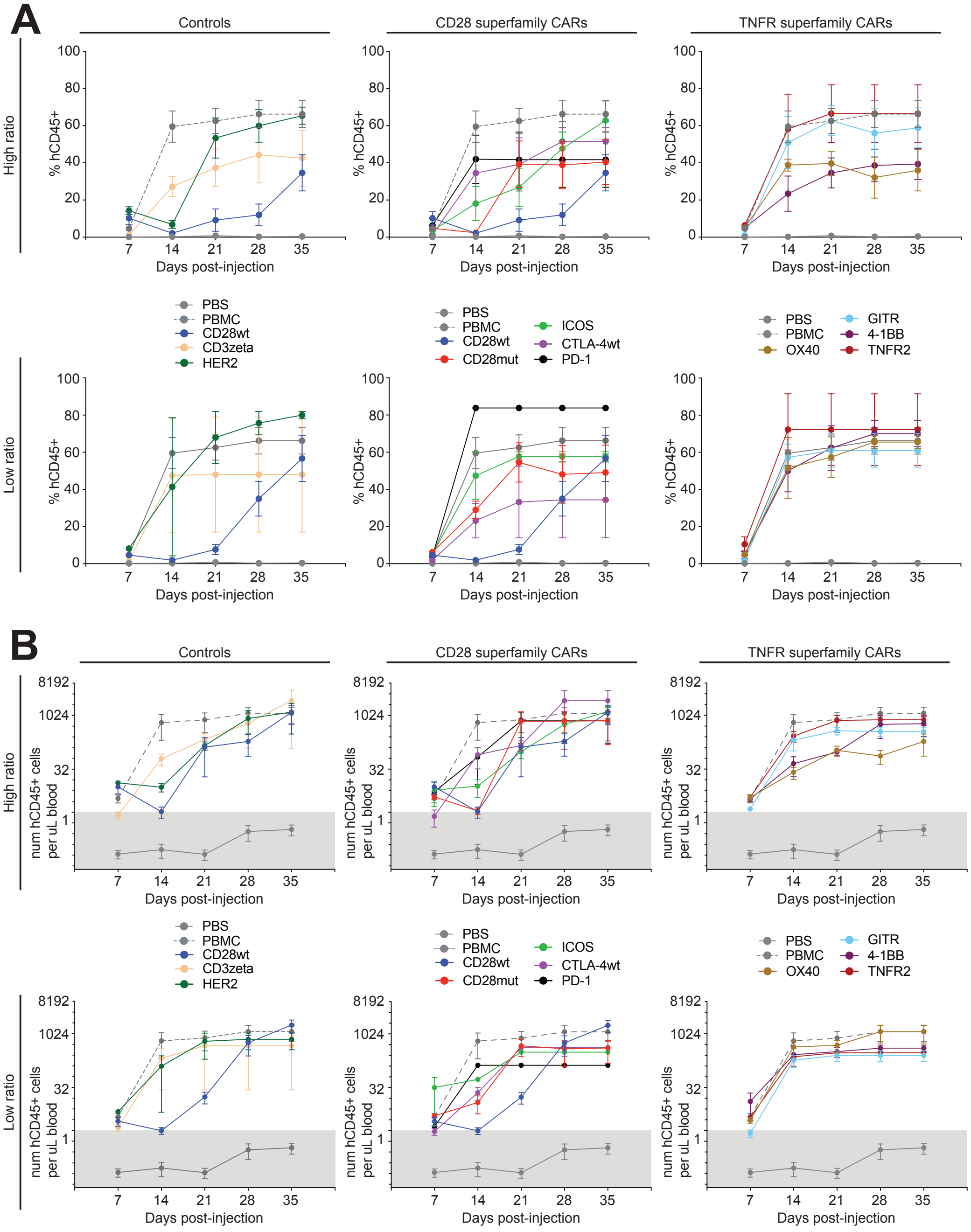


**Supplemental Figure 3. hCD45 engraftment in GVHD model.** Mice were bled weekly and at experimental endpoint. Flow cytometry analysis of blood was performed to determine the blood composition. **(A)** Percent of $hCD45^{+}$ cell engraftment is shown over time, split by Treg:PBMC ratio and CAR signaling domain superfamily. **(B)** Absolute number of $hCD45^{+}$ cells per uL of blood is shown over time. Mean ± SEM.

Supplemental Figure 4

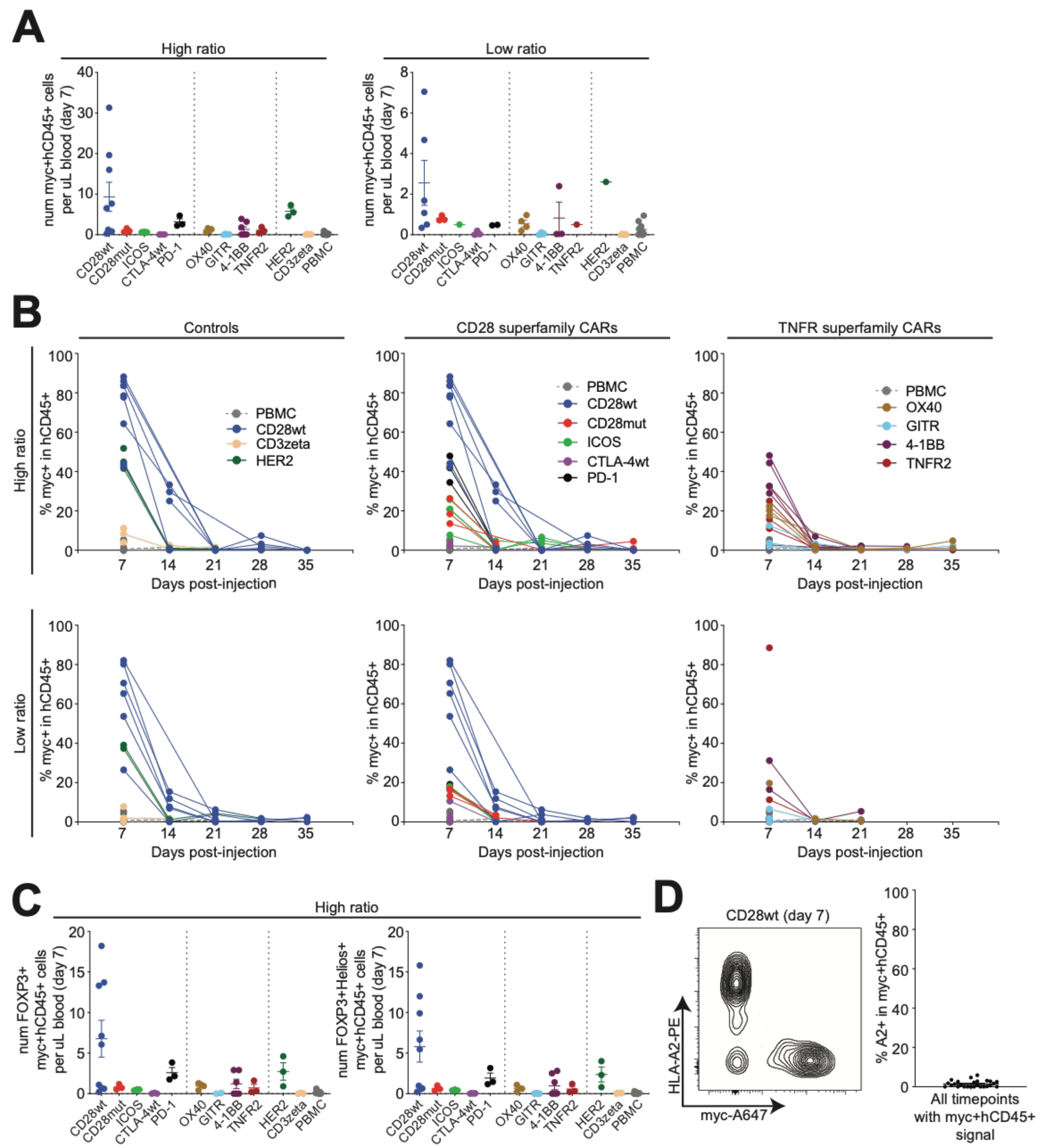


**Supplemental Figure 4. Absolute number of Myc⁺hCD45⁺ cells in GVHD model.** Mice were bled on day 7 as per Figure 2A. **(A)** Proportion of myc⁺ cells in hCD45⁺ subset is shown over time, split by Treg:PBMC ratio and CAR signaling domain superfamily. **(B)** Absolute number of myc⁺hCD45⁺ cells per uL of blood was determined on day 7 post-cell injection. Left: high Treg:PBMC ratio mice are shown. Right: low Treg:PBMC ratio mice are shown. **(C)** Absolute number of FOXP3⁺ myc⁺hCD45⁺ (left) FOXP3⁺Helios⁺myc⁺hCD45⁺ (right) cells per uL of blood was determined on day 7 post-cell injection. High Treg:PBMC ratio is shown. **(D)** Comparison of HLA-A2- and myc-expressing cells. Left: example flow plot of mutually-exclusive HLA-A2 and myc staining. Right: Summary data are shown. Each dot is the average A2⁺ expression of the myc⁺hCD45⁺ population at a single timepoint. Mean ± SEM.

Supplemental Figure 5

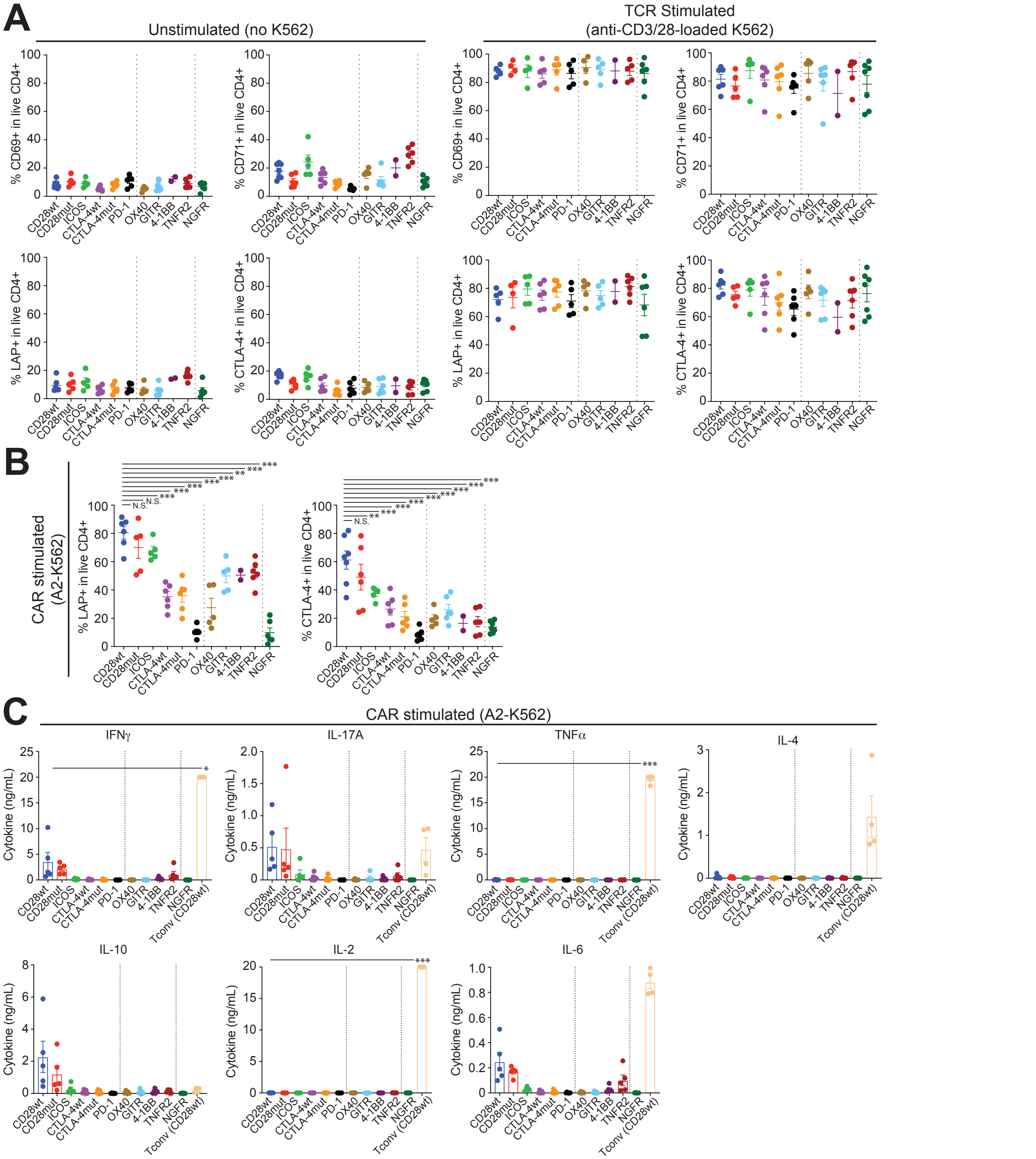


**Supplemental Figure 5. Activation and cytokine production by signaling domain CAR variants on human Tregs. (A-C)** Human Tregs were transduced and expanded as in Figure 1B, rested overnight in low IL-2 conditions, then co-cultured with either no K562 cells (unstimulated), irradiated K562 cells expressing HLA-A2 (CAR-stimulated) or irradiated anti-CD3/CD28-loaded K562 cells expressing CD64 (TCR-stimulated) at a ratio of 1 K562 to 2 CAR Tregs. **(A-B)** After 24 hours, percent CD69, CD71, LAP and CTLA-4 positive in live CD4 cells was determined. Summary data for **(A)** unstimulated and TCR-stimulated cells or **(B)** CAR-stimulated cells are shown. Data are n=2-7 donors, pooled from at least two independent experiments. **(C)** After 72 hours, amounts of the indicated cytokines were determined by cytometric bead array in CAR-stimulated cells. Results for all examined cytokines are expressed as a concentration in the supernatant. n=4-5 from at least two independent experiments. Statistics show one-way ANOVA with Holm-Sidak post-test comparing all constructs to CD28wt Tregs. Mean ± SEM. * $p < 0.05$, ** $p < 0.01$, *** $p < 0.001$. “n.s.” denotes not significant.

Supplemental Figure 6

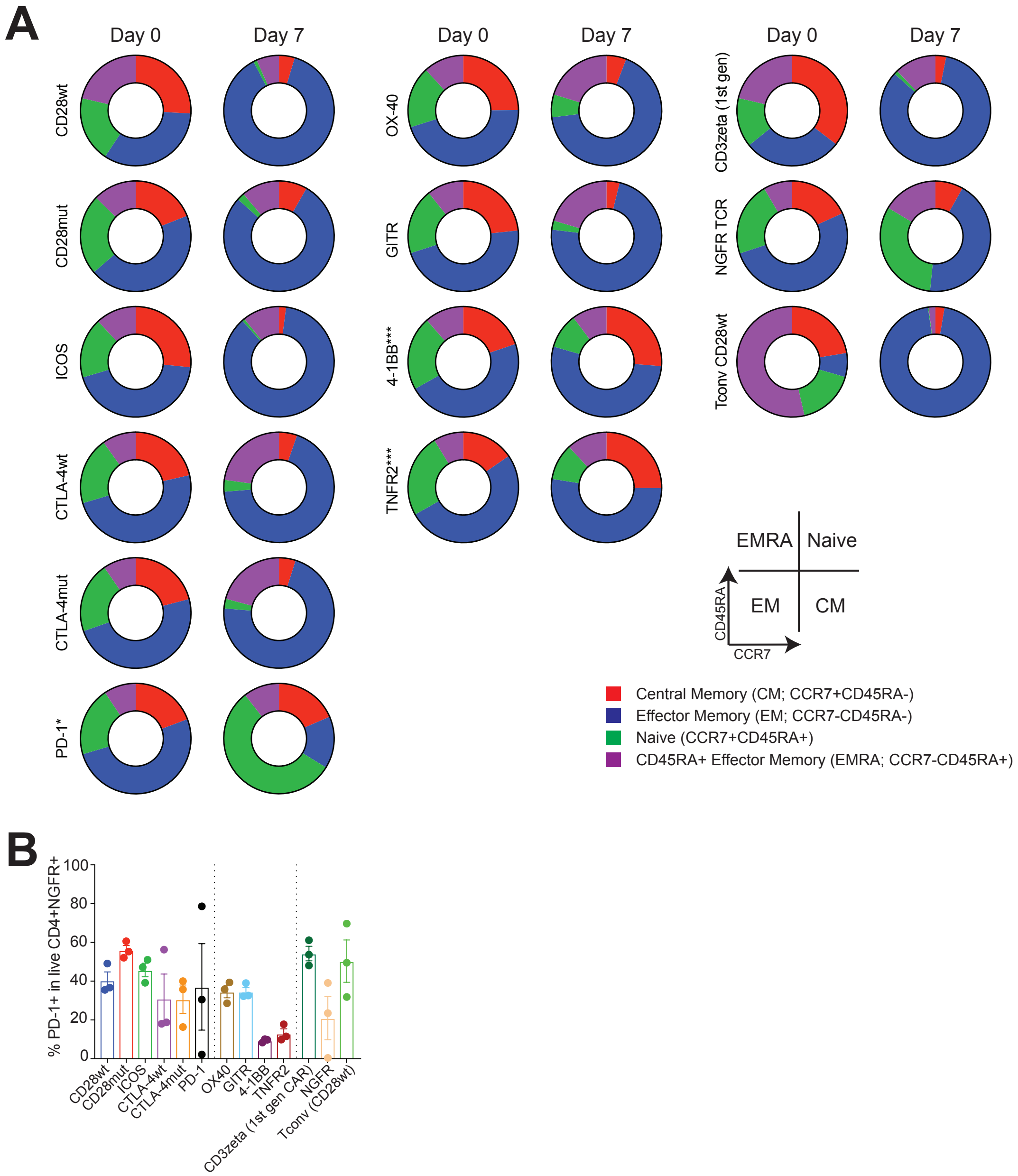


**Supplemental Figure 6. Memory subset and exhaustion of signaling variant CAR Tregs after 7-day CAR-mediated expansion.** Signaling-variant CAR Tregs derived from naïve Tregs were produced as in Figure 5, then co-cultured for 7 days with irradiated K562 cells expressing HLA-A2 at a ratio of 1 K562 cell to 2 CAR Tregs. **(A)** Using flow cytometry analysis of CD45RA and CCR7 expression in live $CD4^{+}NGFR^{+}$ cells, memory subset polarization after CAR stimulation was determined. Average subset values are shown for baseline (day 0) and day 7. Statistics show a one-way ANOVA with Holm-Sidak post-test comparing proportion of central memory cells ($CD45RA^{-}CCR7^{+}$) from CD28wt to all other groups. **(B)** PD-1 expression of $CD4^{+}NGFR^{+}$ cells. Data are from n=3-4 donors, pooled from at least 2 independent experiments. *** $p < 0.001$.

Supplemental Figure 7

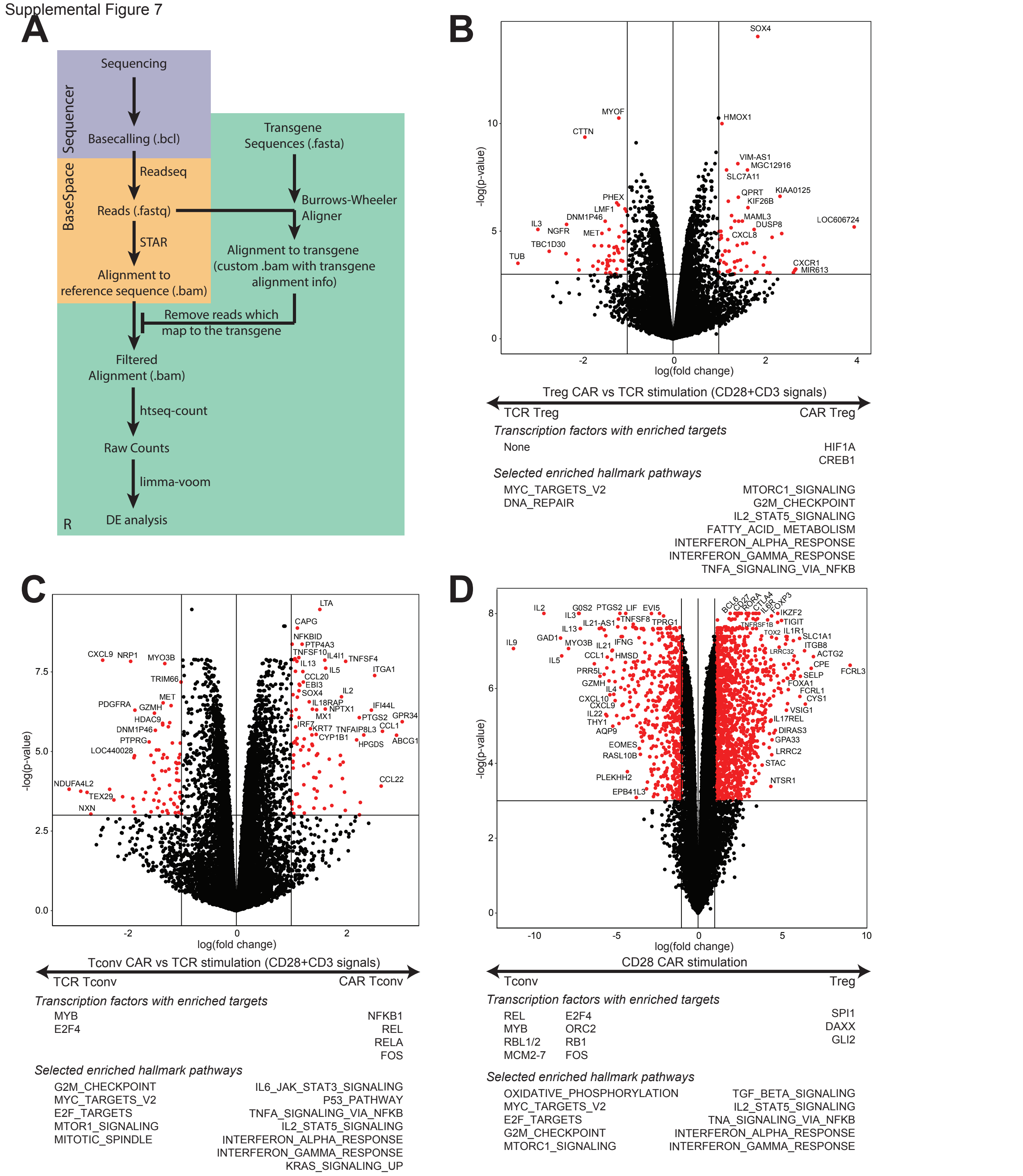


**Supplemental Figure 7. Transcriptome analysis of CAR- and TCR-stimulated Tregs and Tconv.** Purified ΔNGFR/CAR Tregs or Tconv were stimulated with anti-CD3/CD28-dynabeads or HLA-A2-coated beads for 16 hours, then processed for RNA sequencing. **(A)** Schematic of RNA-sequencing pre-processing strategy to remove transgene reads. Complete description of actions performed is written in Methods. To observe direct functional consequences of CAR-stimulation, any sequencing reads that mapped to the CAR transgene or ΔNGFR transduction marker were removed from further differential gene expression analysis. **(B)** Differentially expressed genes between CD28wt-CAR- and TCR-stimulated Tregs. **(C)** Differentially expressed genes between CD28wt-CAR- and TCR-stimulated Tconv. **(D)** Differentially expressed genes between CAR-stimulated Tregs and Tconv expressing a CD28wt-CAR. Full lists, normalized enrichment scores and adjusted p values are found in **Supplemental Table 5-7**.

**Supplemental Table 1. Gene set enrichment analysis of transcription factor targets and hallmark pathways supporting Figure 7A**

| Group: PD-1, TNFR2, CTLA-4mut, CTLA-4wt, CD3zeta | | | Group: CD28wt | | |
|---|---|---|---|---|---|
| **Enriched transcription factor targets** | **NES** | **Adj. p val.** | **Enriched transcription factor targets** | **NES** | **Adj. p val.** |
| CREBBP | -1.92 | 0.0333 | MYB | 2.00 | 0.0192 |
| EP300 | -1.92 | 0.0333 | ORC2 | 1.99 | 0.0213 |
| | | | RB1 | 1.94 | 0.0329 |
| | | | E2F4 | 1.90 | 0.0192 |
| | | | MCM2 | 1.87 | 0.0402 |
| | | | MCM3 | 1.87 | 0.0402 |
| | | | MCM4 | 1.87 | 0.0402 |
| | | | MCM5 | 1.87 | 0.0402 |
| | | | MCM6 | 1.87 | 0.0402 |
| | | | MCM7 | 1.87 | 0.0402 |
| | | | RELA | 1.62 | 0.0402 |
| | | | NFKB1 | 1.45 | 0.0449 |

| Enriched hallmark pathways | NES | Adj. p val. | Enriched hallmark pathways | NES | Adj. p val. |
|---|---|---|---|---|---|
| HALLMARK_INTERFERON_ALPHA_RESPONSE | -2.01 | 0.001 | HALLMARK_MYC_TARGETS_V1 | 3.43 | 0.001 |
| HALLMARK_INTERFERON_GAMMA_RESPONSE | -1.73 | 0.001 | HALLMARK_MYC_TARGETS_V2 | 3.08 | 0.001 |
| HALLMARK_HEME_METABOLISM | -1.60 | 0.004 | HALLMARK_E2F_TARGETS | 2.97 | 0.001 |
| | | | HALLMARK_MTORC1_SIGNALING | 2.62 | 0.001 |
| | | | HALLMARK_UNFOLDED_PROTEIN_RESPONSE | 2.58 | 0.001 |
| | | | HALLMARK_G2M_CHECKPOINT | 2.50 | 0.001 |
| | | | HALLMARK_OXIDATIVE_PHOSPHORYLATION | 2.07 | 0.001 |
| | | | HALLMARK_GLYCOLYSIS | 2.05 | 0.001 |
| | | | HALLMARK_DNA_REPAIR | 2.04 | 0.001 |
| | | | HALLMARK_UV_RESPONSE_UP | 1.86 | 0.001 |
| | | | HALLMARK_ADIPOGENESIS | 1.83 | 0.001 |
| | | | HALLMARK_FATTY_ACID_METABOLISM | 1.48 | 0.022 |

**Supplemental Table 2. Gene set enrichment analysis of transcription factor targets and hallmark pathways supporting Figure 7B**

| Group: 4-1BB, TNFR2 | | | Group: CD28wt | | |
|---|---|---|---|---|---|
| **Enriched transcription factor targets** | **NES** | **Adj. p val.** | **Enriched transcription factor targets** | **NES** | **Adj. p val.** |
| IKBKG | -1.93 | 0.0194 | ORC2 | 1.98 | 0.0190 |
| CREBBP | -1.90 | 0.0190 | MCM2 | 1.83 | 0.0473 |
| EP300 | -1.90 | 0.0190 | MCM3 | 1.83 | 0.0473 |
| | | | MCM4 | 1.83 | 0.0473 |
| | | | MCM5 | 1.83 | 0.0473 |
| | | | MCM6 | 1.83 | 0.0473 |
| | | | MCM7 | 1.83 | 0.0473 |
| | | | E2F4 | 1.71 | 0.0190 |
| | | | MYB | 1.63 | 0.0194 |

| Enriched hallmark pathways | NES | Adj. p val. | Enriched hallmark pathways | NES | Adj. p val. |
|---|---|---|---|---|---|
| HALLMARK_MYC_TARGETS_V2 | -2.76 | 0.003 | HALLMARK_INTERFERON_ALPHA_RESPONSE | 2.20 | 0.003 |
| HALLMARK_MYC_TARGETS_V1 | -2.74 | 0.003 | HALLMARK_INTERFERON_GAMMA_RESPONSE | 2.08 | 0.003 |
| HALLMARK_E2F_TARGETS | -2.60 | 0.003 | HALLMARK_TNFA_SIGNALING_VIA_NFKB | 1.52 | 0.007 |
| HALLMARK_G2M_CHECKPOINT | -2.21 | 0.003 | HALLMARK_HEME_METABOLISM | 1.41 | 0.037 |
| HALLMARK_MTORC1_SIGNALING | -2.08 | 0.003 | | | |
| HALLMARK_UNFOLDED_PROTEIN_RESPONSE | -2.08 | 0.003 | | | |
| HALLMARK_UV_RESPONSE_UP | -1.59 | 0.006 | | | |
| HALLMARK_GLYCOLYSIS | -1.48 | 0.013 | | | |
| HALLMARK_IL2_STAT5_SIGNALING | -1.44 | 0.013 | | | |

**Supplemental Table 3. Gene set enrichment analysis of transcription factor targets and hallmark pathways supporting Figure 7C**

| Group: PD-1 | | | Group: CD3zeta | | |
|---|---|---|---|---|---|
| **Enriched transcription factor targets** | **NES** | **Adj. p val.** | **Enriched transcription factor targets** | **NES** | **Adj. p val.** |
| STAT6 | -1.96 | 0.0325 | MYB | 2.06 | 0.0276 |
| STAT2 | -1.88 | 0.0420 | E2F4 | 1.46 | 0.0430 |
| STAT3 | -1.69 | 0.0420 | | | |

| Enriched hallmark pathways | NES | Adj. p val. | Enriched hallmark pathways | NES | Adj. p val. |
|---|---|---|---|---|---|
| HALLMARK_P53_PATHWAY | -1.42 | 0.0140 | HALLMARK_MYC_TARGETS_V1 | 2.70 | 0.0007 |
| | | | HALLMARK_MYC_TARGETS_V2 | 2.50 | 0.0007 |
| | | | HALLMARK_MTORC1_SIGNALING | 2.32 | 0.0007 |
| | | | HALLMARK_E2F_TARGETS | 2.11 | 0.0007 |
| | | | HALLMARK_UNFOLDED_PROTEIN_RESPONSE | 2.09 | 0.0007 |
| | | | HALLMARK_CHOLESTEROL_HOMEOSTASIS | 2.03 | 0.0007 |
| | | | HALLMARK_TNFA_SIGNALING_VIA_NFKB | 1.90 | 0.0007 |
| | | | HALLMARK_ESTROGEN_RESPONSE_LATE | 1.83 | 0.0007 |
| | | | HALLMARK_G2M_CHECKPOINT | 1.81 | 0.0007 |
| | | | HALLMARK_OXIDATIVE_PHOSPHORYLATION | 1.79 | 0.0007 |
| | | | HALLMARK_GLYCOLYSIS | 1.76 | 0.0007 |
| | | | HALLMARK_FATTY_ACID_METABOLISM | 1.71 | 0.0007 |
| | | | HALLMARK_IL2_STAT5_SIGNALING | 1.60 | 0.0019 |
| | | | HALLMARK_ANDROGEN_RESPONSE | 1.56 | 0.0125 |
| | | | HALLMARK_UV_RESPONSE_UP | 1.43 | 0.0181 |
| | | | HALLMARK_HYPOXIA | 1.43 | 0.0181 |
| | | | HALLMARK_ADIPOGENESIS | 1.41 | 0.0170 |
| | | | HALLMARK_XENOBIOTIC_METABOLISM | 1.41 | 0.0249 |
| | | | HALLMARK_INFLAMMATORY_RESPONSE | 1.38 | 0.0340 |
| | | | HALLMARK_DNA_REPAIR | 1.38 | 0.0340 |

**Supplemental Table 4. Gene set enrichment analysis of transcription factor targets and hallmark pathways supporting Figure 7D**

**Group: CD28mut**

| Enriched transcription factor targets | NES | Adj. p val. |
|---|---|---|
| STAT1 | -1.91 | 0.0336 |

| Enriched hallmark pathways | NES | Adj. p val. |
|---|---|---|
| HALLMARK_WNT_BETA_CATENIN_SIGNALING | -1.75 | 0.0124 |
| HALLMARK_INTERFERON_GAMMA_RESPONSE | -1.67 | 0.0014 |
| HALLMARK_ALLOGRAFT_REJECTION | -1.56 | 0.0064 |
| HALLMARK_INTERFERON_ALPHA_RESPONSE | -1.51 | 0.0208 |

**Group: CD28wt**

| Enriched transcription factor targets | NES | Adj. p val. |
|---|---|---|
| E2F4 | 1.70 | 0.0269 |

| Enriched hallmark pathways | NES | Adj. p val. |
|---|---|---|
| HALLMARK_E2F_TARGETS | 2.52 | 0.0013 |
| HALLMARK_MYC_TARGETS_V1 | 2.45 | 0.0013 |
| HALLMARK_G2M_CHECKPOINT | 2.35 | 0.0013 |
| HALLMARK_MTORC1_SIGNALING | 2.28 | 0.0013 |
| HALLMARK_MYC_TARGETS_V2 | 2.02 | 0.0013 |
| HALLMARK_PROTEIN_SECRETION | 1.96 | 0.0013 |
| HALLMARK_MITOTIC_SPINDLE | 1.78 | 0.0013 |
| HALLMARK_UNFOLDED_PROTEIN_RESPONSE | 1.72 | 0.0030 |
| HALLMARK_FATTY_ACID_METABOLISM | 1.68 | 0.0032 |
| HALLMARK_XENOBIOTIC_METABOLISM | 1.67 | 0.0032 |
| HALLMARK_SPERMATOGENESIS | 1.56 | 0.0209 |
| HALLMARK_ANDROGEN_RESPONSE | 1.54 | 0.0209 |
| HALLMARK_PEROXISOME | 1.54 | 0.0209 |
| HALLMARK_OXIDATIVE_PHOSPHORYLATION | 1.50 | 0.0093 |
| HALLMARK_COMPLEMENT | 1.50 | 0.0148 |
| HALLMARK_ADIPOGENESIS | 1.50 | 0.0129 |
| HALLMARK_GLYCOLYSIS | 1.43 | 0.0239 |

**Supplemental Table 5. Gene set enrichment analysis of transcription factor targets and hallmark pathways supporting Supplemental Figure 7B**

| Group: Treg TCR (CD3+CD28) | | | Group: Treg CAR CD28wt | | |
|---|---|---|---|---|---|
| **Enriched transcription factor targets** | **NES** | **Adj. p val.** | **Enriched transcription factor targets** | **NES** | **Adj. p val.** |
| | | | HIF1A | 1.88 | 0.0216 |
| | | | CREB1 | 1.81 | 0.0216 |
| **Enriched hallmark pathways** | **NES** | **Adj. p val.** | **Enriched hallmark pathways** | **NES** | **Adj. p val.** |
| HALLMARK_MYC_TARGETS_V2 | -2.41 | 0.0019 | HALLMARK_CHOLESTEROL_HOMEOSTASIS | 1.84 | 0.0024 |
| HALLMARK_MYC_TARGETS_V1 | -2.32 | 0.0019 | HALLMARK_MITOTIC_SPINDLE | 1.79 | 0.0019 |
| HALLMARK_DNA_REPAIR | -1.44 | 0.0167 | HALLMARK_HYPOXIA | 1.77 | 0.0019 |
| | | | HALLMARK_INTERFERON_GAMMA_RESPONSE | 1.74 | 0.0019 |
| | | | HALLMARK_IL2_STAT5_SIGNALING | 1.74 | 0.0019 |
| | | | HALLMARK_TNFA_SIGNALING_VIA_NFKB | 1.72 | 0.0019 |
| | | | HALLMARK_SPERMATOGENESIS | 1.67 | 0.0043 |
| | | | HALLMARK_HEME_METABOLISM | 1.65 | 0.0019 |
| | | | HALLMARK_INFLAMMATORY_RESPONSE | 1.64 | 0.0024 |
| | | | HALLMARK_INTERFERON_ALPHA_RESPONSE | 1.62 | 0.0055 |
| | | | HALLMARK_REACTIVE_OXIGEN_SPECIES_PATHWAY | 1.60 | 0.0197 |
| | | | HALLMARK_MTORC1_SIGNALING | 1.60 | 0.0024 |
| | | | HALLMARK_KRAS_SIGNALING_UP | 1.60 | 0.0024 |
| | | | HALLMARK_P53_PATHWAY | 1.56 | 0.0024 |
| | | | HALLMARK_APOPTOSIS | 1.54 | 0.0044 |
| | | | HALLMARK_PROTEIN_SECRETION | 1.50 | 0.0182 |
| | | | HALLMARK_ANDROGEN_RESPONSE | 1.50 | 0.0182 |
| | | | HALLMARK_G2M_CHECKPOINT | 1.44 | 0.0043 |
| | | | HALLMARK_FATTY_ACID_METABOLISM | 1.42 | 0.0192 |
| | | | HALLMARK_ALLOGRAFT_REJECTION | 1.39 | 0.0182 |
| | | | HALLMARK_COMPLEMENT | 1.38 | 0.0245 |
| | | | HALLMARK_XENOBIOTIC_METABOLISM | 1.38 | 0.0254 |

**Supplemental Table 6. Gene set enrichment analysis of transcription factor targets and hallmark pathways supporting Supplemental Figure 7C**

| Group: Tconv TCR (CD3+CD28) | | | Group: Tconv CAR CD28wt | | |
|---|---|---|---|---|---|
| **Enriched transcription factor targets** | **NES** | **Adj. p val.** | **Enriched transcription factor targets** | **NES** | **Adj. p val.** |
| MYB | -1.69 | 0.0124 | REL | 1.97 | 0.0175 |
| E2F4 | -1.59 | 0.0175 | NFKB1 | 1.86 | 0.0124 |
| | | | RELA | 1.84 | 0.0124 |
| | | | FOS | 1.75 | 0.0234 |

| Enriched hallmark pathways | NES | Adj. p val. | Enriched hallmark pathways | NES | Adj. p val. |
|---|---|---|---|---|---|
| HALLMARK_G2M_CHECKPOINT | -2.57 | 0.0010 | HALLMARK_TNFA_SIGNALING_VIA_NFKB | 2.04 | 0.0010 |
| HALLMARK_MYC_TARGETS_V1 | -2.42 | 0.0010 | HALLMARK_INTERFERON_ALPHA_RESPONSE | 2.03 | 0.0010 |
| HALLMARK_E2F_TARGETS | -2.17 | 0.0010 | HALLMARK_INFLAMMATORY_RESPONSE | 1.90 | 0.0010 |
| HALLMARK_MYC_TARGETS_V2 | -1.96 | 0.0010 | HALLMARK_P53_PATHWAY | 1.84 | 0.0010 |
| HALLMARK_MITOTIC_SPINDLE | -1.83 | 0.0010 | HALLMARK_IL2_STAT5_SIGNALING | 1.80 | 0.0010 |
| HALLMARK_UV_RESPONSE_DN | -1.72 | 0.0010 | HALLMARK_INTERFERON_GAMMA_RESPONSE | 1.77 | 0.0010 |
| HALLMARK_UNFOLDED_PROTEIN_RESPONSE | -1.36 | 0.0487 | HALLMARK_XENOBIOTIC_METABOLISM | 1.62 | 0.0026 |
| HALLMARK_MTORC1_SIGNALING | -1.32 | 0.0429 | HALLMARK_REACTIVE_OXIGEN_SPECIES_PATHWAY | 1.50 | 0.0451 |
| | | | HALLMARK_APOPTOSIS | 1.50 | 0.0104 |
| | | | HALLMARK_IL6_JAK_STAT3_SIGNALING | 1.46 | 0.0475 |
| | | | HALLMARK_UV_RESPONSE_UP | 1.46 | 0.0194 |
| | | | HALLMARK_EPITHELIAL_MESENCHYMAL_TRANSITION | 1.44 | 0.0248 |
| | | | HALLMARK_COMPLEMENT | 1.41 | 0.0248 |
| | | | HALLMARK_KRAS_SIGNALING_UP | 1.40 | 0.0328 |
| | | | HALLMARK_APICAL_JUNCTION | 1.37 | 0.0429 |

**Supplemental Table 7. Gene set enrichment analysis of transcription factor targets and hallmark pathways supporting Supplemental Figure 7D**

| Group: Tconv CD28wt | | | Group: Treg CD28wt | | |
|---|---|---|---|---|---|
| **Enriched transcription factor targets** | **NES** | **Adj. p val.** | **Enriched transcription factor targets** | **NES** | **Adj. p val.** |
| REL | -2.34 | 0.0082 | SPI1 | 2.03 | 0.0082 |
| MYB | -2.07 | 0.0082 | DAXX | 1.93 | 0.0230 |
| RBL1 | -2.04 | 0.0230 | GLI2 | 1.85 | 0.0331 |
| RBL2 | -2.04 | 0.0230 | | | |
| MCM2 | -1.93 | 0.0230 | | | |
| MCM3 | -1.93 | 0.0230 | | | |
| MCM4 | -1.93 | 0.0230 | | | |
| MCM5 | -1.93 | 0.0230 | | | |
| MCM6 | -1.93 | 0.0230 | | | |
| MCM7 | -1.93 | 0.0230 | | | |
| E2F4 | -1.90 | 0.0082 | | | |
| RB1 | -1.83 | 0.0294 | | | |
| ORC2 | -1.82 | 0.0331 | | | |
| FOS | -1.71 | 0.0309 | | | |
| **Enriched hallmark pathways** | **NES** | **Adj. p val.** | **Enriched hallmark pathways** | **NES** | **Adj. p val.** |
| HALLMARK_MYC_TARGETS_V1 | -3.71 | 0.0011 | HALLMARK_INTERFERON_ALPHA_RESPONSE | 2.06 | 0.0011 |
| HALLMARK_MYC_TARGETS_V2 | -3.38 | 0.0011 | HALLMARK_CHOLESTEROL_HOMEOSTASIS | 1.88 | 0.0013 |
| HALLMARK_E2F_TARGETS | -2.88 | 0.0011 | HALLMARK_HYPOXIA | 1.87 | 0.0013 |
| HALLMARK_UNFOLDED_PROTEIN_RESPONSE | -2.41 | 0.0011 | HALLMARK_HEME_METABOLISM | 1.80 | 0.0013 |
| HALLMARK_DNA_REPAIR | -2.18 | 0.0011 | HALLMARK_APOPTOSIS | 1.80 | 0.0011 |
| HALLMARK_G2M_CHECKPOINT | -2.16 | 0.0011 | HALLMARK_INTERFERON_GAMMA_RESPONSE | 1.80 | 0.0013 |
| HALLMARK_OXIDATIVE_PHOSPHORYLATION | -1.95 | 0.0011 | HALLMARK_UV_RESPONSE_DN | 1.75 | 0.0013 |
| HALLMARK_MTORC1_SIGNALING | -1.78 | 0.0011 | HALLMARK_COMPLEMENT | 1.75 | 0.0016 |
| HALLMARK_UV_RESPONSE_UP | -1.50 | 0.0079 | HALLMARK_IL6_JAK_STAT3_SIGNALING | 1.70 | 0.0063 |
| | | | HALLMARK_TNFA_SIGNALING_VIA_NFKB | 1.69 | 0.0013 |
| | | | HALLMARK_P53_PATHWAY | 1.65 | 0.0013 |
| | | | HALLMARK_MITOTIC_SPINDLE | 1.57 | 0.0016 |
| | | | HALLMARK_IL2_STAT5_SIGNALING | 1.57 | 0.0016 |
| | | | HALLMARK_MYOGENESIS | 1.55 | 0.0074 |

| Enriched hallmark pathways | NES | Adj. p val. | Enriched hallmark pathways | NES | Adj. p val. |
|---|---|---|---|---|---|
| | | | HALLMARK_APICAL_JUNCTION | 1.53 | 0.0063 |
| | | | HALLMARK_EPITHELIAL_MESENCHYMAL_TRANSITION | 1.49 | 0.0114 |
| | | | HALLMARK_BILE_ACID_METABOLISM | 1.49 | 0.0244 |
| | | | HALLMARK_TGF_BETA_SIGNALING | 1.49 | 0.0385 |
| | | | HALLMARK_ALLOGRAFT_REJECTION | 1.48 | 0.0087 |
| | | | HALLMARK_PROTEIN_SECRETION | 1.47 | 0.0241 |
| | | | HALLMARK_KRAS_SIGNALING_DN | 1.43 | 0.0337 |
| | | | HALLMARK_KRAS_SIGNALING_UP | 1.40 | 0.0275 |
